# Activation of amino acid metabolic program in response to impaired glycolysis in cardiac HIF1 deficient mice

**DOI:** 10.1101/2020.05.23.111674

**Authors:** Ivan Menendez-Montes, Beatriz Escobar, Manuel J. Gomez, Teresa Albendea-Gomez, Beatriz Palacios, Elena Bonzon, Ana Vanessa Alonso, Alessia Ferrarini, Luis Jesus Jimenez-Borreguero, Jesus Vázquez, Silvia Martin-Puig

## Abstract

Hypoxia is an important environmental cue in heart development. Despite of extensive characterization of gain and loss of function models, there is disagreement about the impact of HIF1α elimination in cardiac tissue. Here, we used a new conditional knock out of *Hif1a* in NKX2.5 cardiac progenitors to assess the morphological and functional consequences of HIF1α loss in the developing heart. By combining histology, electron microscopy and high-throughout genomics, proteomics and metabolomics, we found that deletion of Hif1a leads to impaired embryonic glycolysis without influencing cardiomyocyte proliferation and results in an increased mitochondrial number, activation of a transient amino acid response and upregulation of HIF2α and ATF4 by E12.5. *Hif1a* mutants display normal fatty acid oxidation metabolic profile and do not show any sign of cardiac dysfunction in the adulthood. Our results demonstrate that HIF1 signaling is dispensable for heart development and reveal the metabolic flexibility of the embryonic myocardium, opening the potential application of alternative energy sources as therapeutic interventions during ischemic events.

## INTRODUCTION

The heart is the first organ to form during development, as it is essential to deliver oxygen and nutrients to embryonic tissues from early stages. Different subsets of cardiac progenitors proliferate, migrate and differentiate into the diverse cell types that form the mature heart (Martin-Puig et al., 2008; Watanabe and Buckingham, 2010). Despite of the existence of several types of cardiovascular precursors, NKX2.5 progenitors give rise to the majority of cardiac cells, contributing to the three main heart layers: epicardium, myocardium and endocardium (Moses et al., 2001). However, cardiogenesis can result in malformations, and congenital heart defects occur in 1% of live births. Several factors have been involved in developmental cardiac failure, and among them hypoxia, or low oxygen tensions, has been previously described as an environmental factor associated with cardiac defects, including metabolic and vascular alterations (Cerychova and Pavlinkova, 2018; Nanka et al., 2008), during pregnancy. Hypoxia-Inducible Factors (HIFs) are known to mediate a well-characterized transcriptional response to hypoxia. HIFs heterodimers are formed by a constitutively expressed β subunit (HIFβ or ARNT) and an oxygen-regulated α subunit, with three different isoforms (1α, 2α and 3α) (Kaelin and Ratcliffe, 2008). Under normoxic conditions, the oxygen sensors Prolyl Hydroxylases (PHDs) hydroxylate HIFα in specific proline residues (Jiang et al., 1997). This modification is a recognition mark for the von Hippel Lindau/E3 ubiquitin ligase complex, that polyubiquitinates and drives α subunits to proteasomal degradation. In hypoxic conditions, α subunits evade degradation due to the inhibited PHD activity, dimerize with β subunits and mediate the adaptive response to hypoxia by activating the transcriptions of their targets genes (Pouyssegur et al., 2006).

In addition to the existence of hypoxic areas during heart development (Lee et al., 2001), it has been described that chronic exposure of pregnant females to 8% oxygen causes embryonic myocardial thinning, epicardial detachment and death (Ream et al., 2008) and global deletion of different components of the hypoxia pathway display cardiovascular phenotypes (Dunwoodie, 2009). Moreover, genetic-based overactivation of HIF signaling by inactivation of Vhl gene in cardiac progenitors using different drivers (Mlc2vCre, Nkx2.5Cre) causes morphological, metabolic and functional cardiac alterations that results in embryonic lethality (Lei et al., 2008; Menendez-Montes et al., 2016). On the other hand, loss of function models based on conditional deletion of *Hif1a* gene in cardiac progenitor populations also causes several cardiac alterations during embryogenesis (Guimarães-Camboa et al., 2015; Huang et al., 2004; J Krishnan et al., 2008), indicating that a controlled balance in the oxygen levels and hypoxia signaling is required for proper cardiac development. However, there are important phenotypic discrepancies between the different published loss of function models. On one hand, the use of cardiac-specific Mlc2vCre driver in combination with a *Hif1a* floxed mice does not affect embryonic survival but causes cardiac hypertrophy with reduced cardiac function in the adulthood, together with decreased glycolysis and ATP and lactate levels (Huang et al., 2004). On the other hand, when a null *Hif1a* allele in germline is used in combination with a *Hif1a* floxed allele and the Mlc2vCre driver, mutant embryos show several cardiac alterations and increased cardiomyocyte proliferation, with associated embryonic lethality by E12.0 (J Krishnan et al., 2008). A more recent study also uses a combination of null and floxed *Hif1a* alleles, this time under the control of Nkx2.5Cre driver, allowing for more homogeneous and efficient recombination than Mlc2vCre (Guimarães-Camboa et al., 2015). In this case, *Hif1a* deletion seems to have opposite effects, with activation of cell stress pathways that inhibit cardiomyocyte proliferation and lead to myocardial thinning, resulting in embryonic lethality by E15.5. Considering the importance of hypoxia pathway in early hematopoiesis, placentation and vascular development, the potential secondary effect of using *Hif1a*-null alleles on cardiac development cannot be ruled out when interpreting data from some of these mutants. Therefore, while it is clear that sustained HIF1 signaling in the embryonic heart is detrimental for proper cardiac development, there is still disagreement and open debate about the impact of *Hif1a* loss during cardiogenesis and subsequent effects on cardiac function.

It has been widely demonstrated that HIF1 signaling is an important regulator of cellular metabolism, both in physiological and pathological contexts. In addition to glycolytic activation (Majmundar et al., 2010), HIF1 reduces mitochondrial metabolism by repressing pyruvate entry into the mitochondria (Kim et al., 2006) and by promoting COX4 isoform switch from COX4-1 to COX4-2 (Fukuda et al., 2007). Moreover, HIF1 can also limit oxidative metabolism through an inhibitory role on mitochondrial biogenesis (Zhang et al., 2007). Several *in vitro* studies have demonstrated that the embryonic heart relies on glycolysis for energy supply (Chung et al., 2011, 2010), in contrast with the adult heart that sustains most of ATP production through mitochondrial oxidation of fatty acids (FA) (Lopaschuk y Jaswal, 2010). This metabolic switch is coincident with the change in oxygen levels after birth (Puente et al., 2014). However, a recent report from our group has shown that an earlier metabolic shift towards fatty acid oxidation occurs during development at around E14.5, through a mechanism dependent of a decrease in HIF1 signaling in the embryonic myocardium (Menendez-Montes et al., 2016). Despite of the importance of glucose and fatty acids as cardiac energy sources, amino acids can also be used as bioenergetics fuel. Hence, amino acids have the capacity to enter the Krebs cycle at different levels, a phenomenon known as anaplerosis, and to replenish metabolic intermediates that warrant both NADH/FADH_2_ and building blocks production that enable the cells to continue growing under amino acid metabolism. The importance of amino acids as catabolic substrates has been described in several contexts, such as tumor growth (Yue et al., 2017), pulmonary hypertension (Piao et al., 2013) or limited oxygen supply conditions (Bing et al., 1954; Julia et al., 1990). However, the ability of the embryonic heart to catabolize amino acids remains unexplored.

Here, we describe that Hif1α loss during cardiogenesis blunts glycolysis and drives a compensatory metabolic adaptation based on transient activation of amino acid transport and catabolism associated with increased ATF4 and HIF2α levels to maintain energy production prior to the establishment of FA-based metabolism by E14.5. Our results demonstrate that HIF1 signaling is dispensable for cardiogenesis and describe a new role of amino acid metabolism during heart development, opening future research horizons towards studying the ability of the heart to use amino acids as an alternative energy fuel and biosynthetic precursor source in different pathophysiological contexts.

## RESULTS

### HIF1 signaling in cardiac progenitors is dispensable for cardiac development

HIF1α is expressed in the developing myocardium, with a temporal dynamics along midgestation (Guimarães-Camboa et al., 2015; Krishnan et al., 2008; Menendez-Montes et al, 2016). We and others have described the heterogeneous regional distribution of HIF1 signaling with high HIF1α levels in the compact myocardium in contrast with low expression in the trabeculae (Guimaraes Camboa et al. 2015; Menendez-Montes et al., 2016). HIF1 has been proposed to regulate cardiomyocyte proliferation during cardiogenesis. However, opposing results have been reported depending on the Cre line used for recombination and the utilization of one global null allele or two floxed alleles of *Hif1α* (Guimarães-Camboa et al., 2015; Krishnan et al., 2008; Huang et al. 2004). To investigate the role of HIF1 in heart development we generated a cardiac-specific loss of function model using two floxed alleles of *Hif1a* gene in combination with the cardiac progenitor-specific Cre recombinase driver Nkx2.5Cre (*Hif1a*^flox/flox^/*Nkx2.5*^Cre/+^, from here on *Hif1a*/*Nkx2.5*). Cre-mediated recombination was analyzed by agarose gel electrophoresis of cardiac and non-cardiac tissue at E12.5 (Fig. S1A). A 400bp product, corresponding to the processed *Hif1a* allele was only obtained in cardiac tissue in the presence of Cre recombinase activity, indicating that the deletion is specific in cardiac tissue and there is not ectopic recombination. To confirm the deletion efficiency, we analyzed by qPCR at E14.5 the expression of several genes involved in the HIF1 pathway (Fig. S1B). Expression of *Hif1a* itself was quantified by amplification inside and outside of the floxed region. Despite of an increase in *Hif1a* mRNA on the outside region, probably caused by compensatory mechanisms, specific lack of amplification inside the floxed region confirmed efficient recombination. In addition, Phd3, whose expression is dependent on HIF1, showed decreased expression in the *Hif1a*/*Nkx2.5* mutants. However, the expression of other HIF1 targets, like *Vegfa*, or additional elements of the hypoxia pathway like *Hif2a*, *Hif1b* or *Vhl* remained largely unchanged. We also determined HIF1α protein distribution and abundance within the tissue by immunostaining in mutant and control littermates by E12.5 (Fig. S1C). *Hif1a* loss in the mutant embryos resulted in reduced HIF1α staining, with a displacement of the HIF1α channel intensity curve to lower fluorescence intensities (Fig. S1D). It is remarkable that the deletion efficiency detected by qPCR is stronger than the one observed by immunostaining. This is probably because *Hif1a* mRNA after recombination is sufficiently stable to be translated, although the resulting protein is not functional as it lacks the DNA-binding domain located in the N-terminal region of HIF1α. The efficient deletion was further confirmed by Western-Blot of *Hif1a*-deficient hearts at E12.5 (Fig. 1A). The lower weight of HIF1α protein band confirms the presence of a truncated protein in the mutant embryos that is recognized by the HIF1 antibody raised against the C-terminal domain.

**Figure 1.**
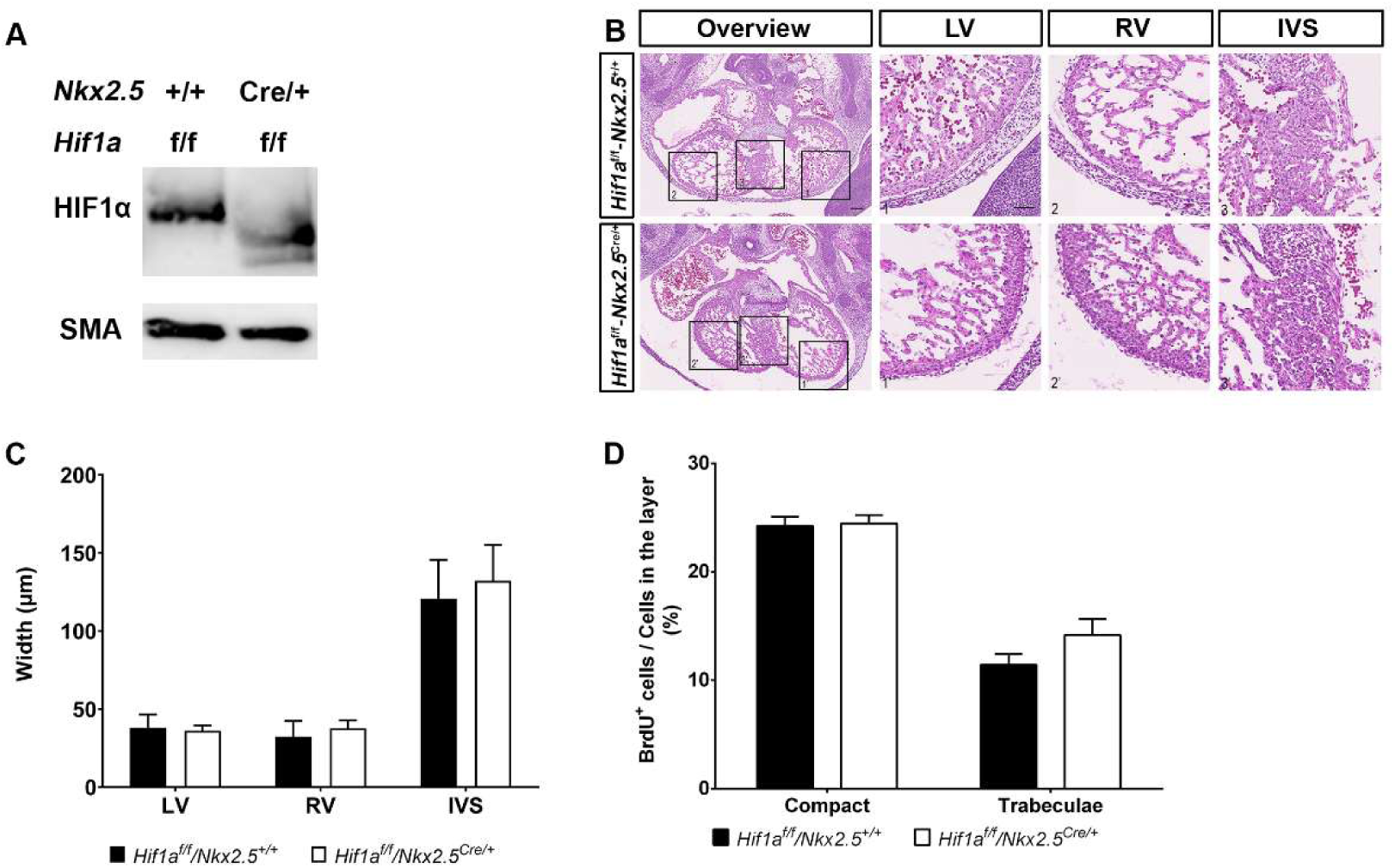
Embryonic phenotype of *Hif1a*-deficient embryos at E12.5. **A)** Representative immunoblot against HIF1α (up) and smooth muscle actin (down) in heart lysates of control (*Hif1a*^f/f^/*Nkx2.5*^+/+^) and *Hif1a*/*Nkx2.5* (*Hif1a*^f/f^/*Nkx2.5*^Cre/+^) embryos at E12.5. **B)** E12.5 control (up) and mutant (down) heart sections stained with hematoxylin and eosin (HE). Scale bars represent 100µm (overview) and 20µm (insets). **C)** HE quantification of ventricular walls and inter-ventricular septum width in E12.5 control (black bars, n=4) and mutant (white bars, n=3) embryos. **D)** Quantification of BrdU immunostaining, represented as percentage of BrdU^+^ cells in the compact myocardium and trabeculae of E12.5 control (black) and *Hif1a*/*Nkx2.5* mutant (white) embryos (n=3). In all graphs, bars represent mean±SEM, Student’s t test, * pvalue<0.05.

*Hif1a*/*Nkx2.5* mutants were viable and recovered in the expected Mendelian proportions from E14.5 to weaning (Table 1). Histological analysis of control and *Hif1a*/*Nkx2.5* mutant hearts at E12.5 (Fig. 1B), did not reveal differences between genotypes in terms of ventricular wall thickness (Fig. 1C) or chamber sphericity (data not shown). Proliferation analysis by BrdU staining at E12.5 proved comparable proliferation index between control and *Hif1a*-deficient hearts (Fig. 1D). Similar results were obtained at E14.5, when *Hif1a*-deficient embryos did not show morphological alterations (Fig. S2A-B) nor differences between genotypes in terms of cell size (Fig. S2C) and proliferation by means of BrdU staining (Fig. S2D). These results indicate that the lack of *Hif1a* in Nkx2.5 cardiac progenitors does not influence cardiomyocyte proliferation and suggest that HIF1 signaling is dispensable for proper cardiac development.

**Table 1.** Embryo recovery analysis of *Hif1a*/*Nkx2.5* mutants. For each stage, table shows the number of mutant embryos recovered, the total number of embryos/pups collected and the number of litters analyzed. The percentage of recovered mutants and the expected recovery percentage (25% in all cases) were compared by the Wilcoxon signed rank test.

| Stage | *Hif1a<sup>f/f</sup>/Nkx2.5<sup>Cre/+</sup>* | Total | Litters | Observed % | Expected % | p-value |
| --- | --- | --- | --- | --- | --- | --- |
| E14.5 | 20 | 90 | 13 | 24.4±4.9 | 25 | 0.905 |
| Weaning | 10 | 38 | 6 | 28.6±5.8 | 25 | 0.787 |

### Developmental loss of *Hif1a* does not influence adult cardiac function or morphology

Albeit deletion of *Hif1a* did not hamper embryonic ventricular chamber formation, we wondered whether mutant mice might develop heart alterations in the adulthood. Histology analysis by Hematoxylin-Eosin staining at five months of age did not indicate evident changes in cardiac morphology of *Hif1a*-deficient hearts relative to control animals (Fig. 2A). Moreover, Mason’s Trichrome staining analysis excluded the presence of fibrotic areas in any of the genotypes (Fig. 2B). Nevertheless, the lack of macroscopic malformations does not rule out that cardiac performance could be affected. To determine if embryonic deletion of *Hif1a* influences cardiac function during adulthood, we performed echocardiography in five months-old control and mutant mice. Both 2D and M mode analysis (Fig. 2C) and the quantification of several echocardiographic parameters (Fig. S3A-D) confirmed the absence of anatomical alterations. Furthermore, conserved cardiac function in *Hif1a*/*Nkx2.5* mutant versus control mice was demonstrated by means of Ejection Fraction (EF) and Fractional Shortening (FS) (Fig. 2D). On the other hand, electrocardiographic analysis showed normal PR and QRS segment length in *Hif1a*/*Nkx2.5* mice (data not shown), ruling out the existence of conduction or coupling defects.

**Figure 2.**
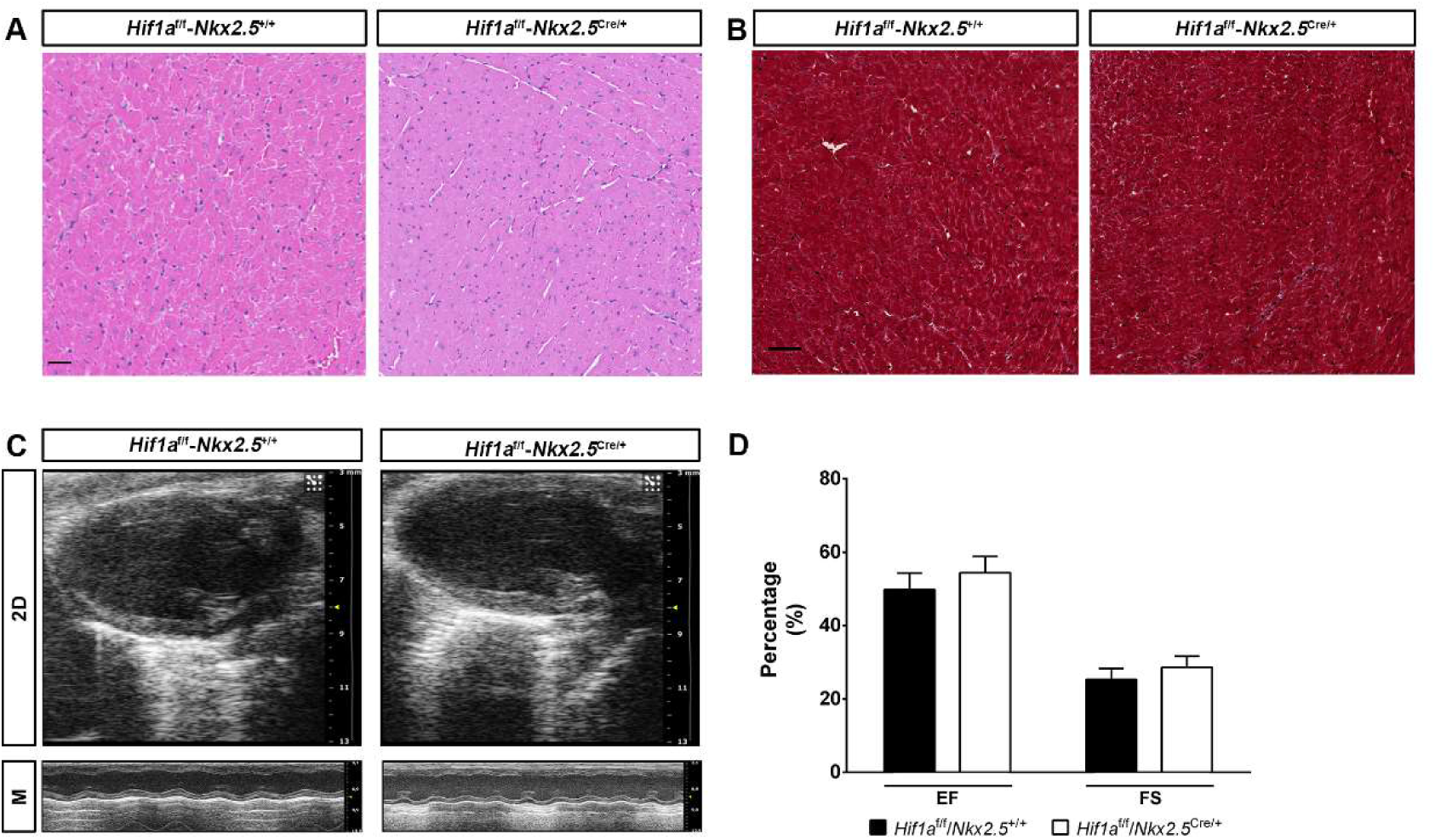
Cardiac morphology and function in adult *Hif1a*/*Nkx2.5* mutants. HE **(A)** and Mason’s Trichromic **(B)** Stainings of the left ventricle from 5 months-old control (*Hif1a*^f/f^/*Nkx2*^*r*+/+^) and *Hif1a*/*Nkx2.5* mutants (*Hif1a*^f/f^/*Nkx2.5*^Cre/+^). Scale bar represents 50µm. **C)** Echocardiography analysis of 5 months-old control and *Hif1a*-deficient mutants in 2D mode (up) and M mode (down). **D)** Quantification of Ejection Fraction (EF) and Fractional Shortening (FS) in control (black bars, n=9) and *Hif1a*/*Nkx2.5* mutants (white bars, n=11) by 5 months of age. Bars represent mean±SEM, Student’s t test. Scale bars represent 50µm.

These results indicate that active HIF1 signaling during heart development is not required for proper cardiac morphogenesis or normal heart function in the adulthood.

### Cardiac deletion of *Hif1a* prevents the expression of glycolytic enzymes in the compact myocardium

To investigate adaptive mechanisms operating upon *Hif1a* loss to allow normal cardiac development, we performed massive expression analysis by RNASeq of E12.5 ventricular tissue from control and *Hif1a*/*Nkx2.5* mutant embryos. Subsequent bioinformatics analysis identified 14406 genes being expressed. Among them, 201 genes showed differential expression: 118 were downregulated and 83 were upregulated in mutant hearts relative to control hearts, representing positively and negatively-regulated targets, in direct or indirect fashion, in the absence of functional HIF1 signaling (Table S1). GO enrichment analysis revealed significant alterations in metabolic processes, such as nucleotide/nucleoside and monocarboxylic acid metabolism, amino acid and, especially, carbohydrate metabolism (Table S2, Fig. 3A). We have previously described the existence of metabolic territories in the embryonic myocardium with an enhanced glycolytic signature in the compact myocardium by E12.5 (Menendez-Montes et al 2016). Here we found that glucose transporter 1 (GLUT1) protein levels were significantly reduced in the compact myocardium of the *Hif1a*/*Nkx2.5* mutant embryos by E12.5 (Fig. 3B). The inhibition of the glycolytic program in the *Hif1a*/*Nkx2.5* single mutant was further confirmed by mRNA expression analysis of the critical enzymes *Glut1*, *Pdk1* and *Ldha* by *in situ* hybridization at E12.5 and E14.5 in control and *Hif1a*/*Nkx2.5* mutants (Fig. 3C). Results showed strong inhibition of glycolytic gene expression in the compact myocardium of *Hif1a*-deficient hearts at both stages. This sustained glycolytic inhibition at E14.5 was validated by qPCR (Fig. 3D), including the downregulation of *Slc16a3*, responsible of the transport of monocarboxylic acids, such as lactate, across mitochondrial membrane. NKX2.5 cardiac progenitors contribute to different cardiac layers including myocardium, epicardium and endocardium.

**Figure 3.**
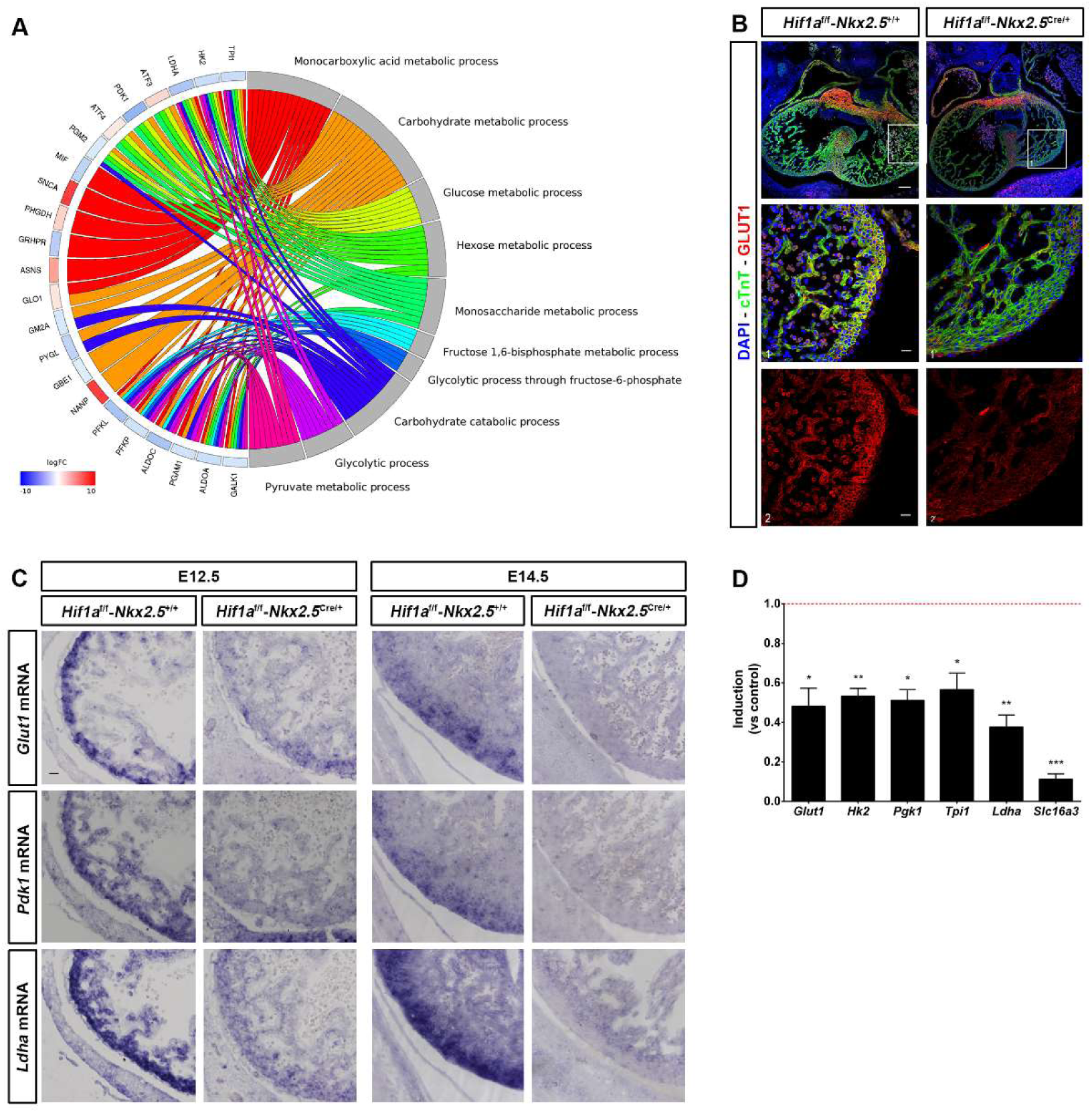
Glycolytic metabolism alterations in *Hif1a*/*Nkx2.5* embryos. **A)** Circular plot representing logFC value for genes detected as differentially expressed in mutant (*Hif1a*^f/f^/*Nkx2.5*^Cre/+^) embryos, relative to control (*Hif1a*^f/f^/*Nkx2.5*^+/+^), at E12.5, associated to a selection of Gene Ontology (GO) terms related to carbohydrate metabolism. Functional terms were detected with GOrilla by comparison against the Biological Process component of the GO database. All terms were enriched with P value < 0.001. logFC values are color coded: red colour denotes higher expression in mutant samples. **B)** Representative GLUT1 immunofluorescence on E12.5 heart sections of control (left) and *Hif1a*/*Nkx2.5* mutants (right). Nuclei shown in blue, Troponin T in green and GLUT1 in red. Insets show left ventricle. Scale bars, 100µm and 20µm in insets. **C)** In situ hybridization of *Glut1* (top), *Pdk1 (middle) and *Ldha* (bottom) in control and *Hif1a*-mutant right ventricle* at E12.5 (left) and E14.5 (right). Scale bar, 20µm **D)** RT-qPCR analysis of glycolytic genes in E14.5 *Hif1a*-mutant ventricles. Bars (mean±SEM, n=3) represent fold induction relative to baseline expression in littermate controls (red line). Student’s t test. *pvalue<0.05; **0.005<pvalue<0.01, ***pvalue<0.005.

To determine if glycolytic inhibition was associated with *Hif1a* loss in the myocardial layer, we specifically deleted *Hif1a* in cardiomyocytes using *cTnT*-Cre (Jiao et al., 2003) (*Hif1a*^flox/flox^/*cTnT*^Cre/+^, hereon *Hif1a*/*cTnT*). *Hif1a/cTnT* mutants also showed reduced HIF1α levels by E14.5 (Fig. S4A) compared with control littermates. Similar to *Hif1a*/*Nkx2.5* mutants, deletion of *Hif1a* in cardiomyocytes did not cause cardiac developmental defects by E14.5 (Fig. S4B) even in the absence of effective glycolysis, as confirmed by *in situ* hybridization (Fig. S4C) and RT-qPCR (Fig.S4D) of key glycolytic pathway members.

Taken together, these results confirm our previous findings that HIF1 signaling controls the expression of glycolytic genes in the embryonic heart and indicate that an active glycolytic program in the compact myocardium is not essential for proper cardiogenesis.

### Lipid metabolism is preserved in cardiac *Hif1a* mutant mice

We have previously reported that sustained HIF1 signaling in the embryonic myocardium results in severe alterations of mitochondrial amount and function (Menendez-Montes et al. 2016). To evaluate the bioenergetics adaptations in response to the lack of cardiac HIF1 activation we investigated mitochondrial network and activity in *Hif1a*/*Nkx2.5* mutants. Analysis and quantification of ventricular ultrastructure by TEM at E12.5 indicated a moderate increase in mitochondrial content in *Hif1a*-deficient embryos compared with control littermates. Images also confirmed our previously reported observation that mitochondrial number is higher in the trabecular layer than in the compact myocardial layer (Menendez-Montes et al., 2016), both in control and *Hif1a* mutant hearts (Fig 4A). Enriched mitochondrial content in *Hif1a*/*Nkx2.5* mutants correlated with reduced expression levels of HIF1 target genes involved in mitophagy like *Nix* and *Bnip3* (Zhang et al., 2008) or genes that negatively regulate mitochondrial biogenesis through Myc transcriptional repression like *Mxi1* (max interacting protein 1) (Fig 4B). Furthermore, we observed a significant enriched expression of genes related to oxidative phosphorylation determined by Gene Set Enrichment Analysis (GSEA) (Table S3, Fig 4C).

**Figure 4.**
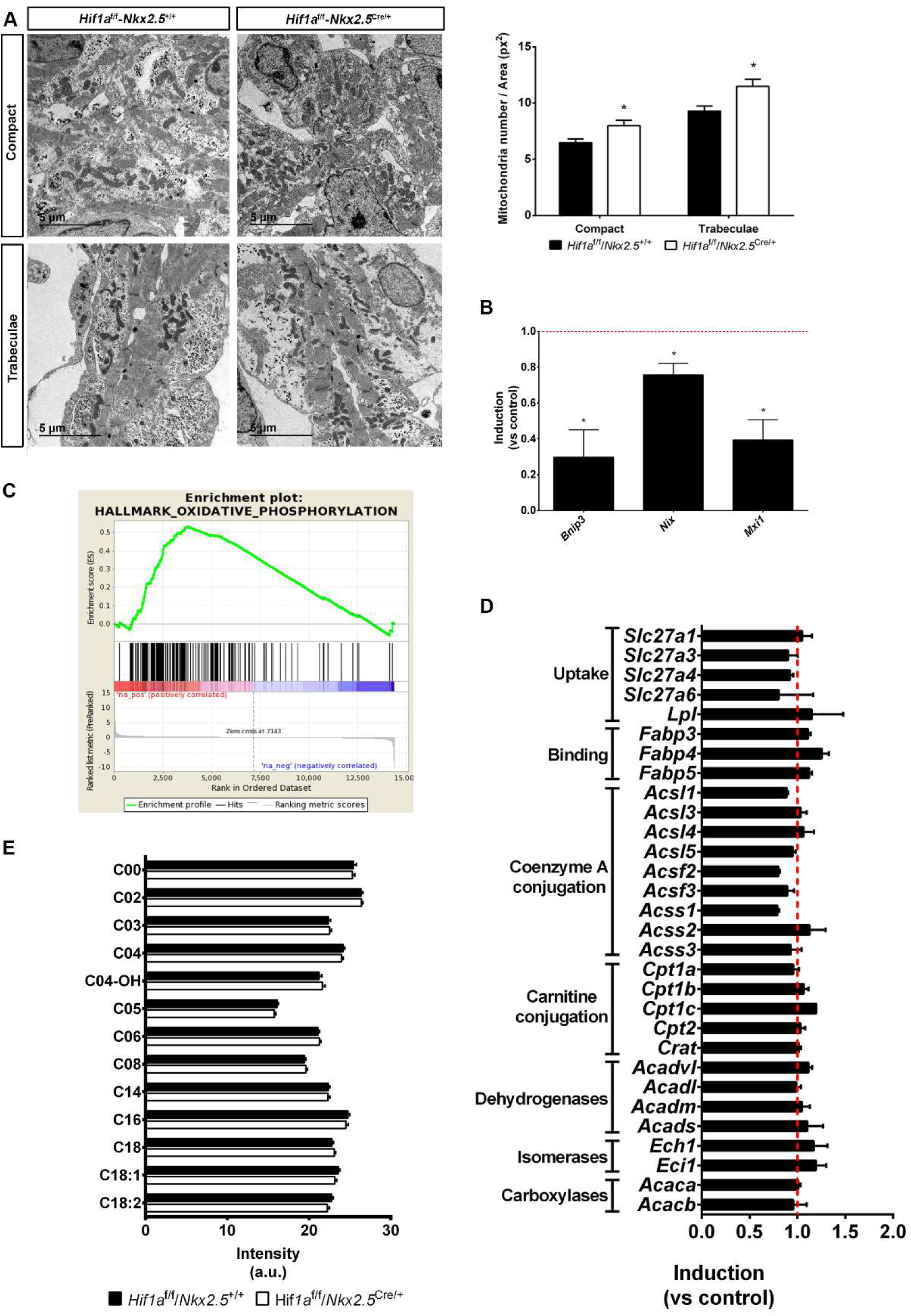
Mitochondrial content and lipid metabolism in *Hif1a*/*Nkx2.5* mutants at E12.5. **A)** Transmission electron micrographs of ventricular tissue from a representative E12.5 control embryo (*Hif1a*^*f/f*^/*Nkx2.5*^*+/+*^, left) and a mutant littermate (*Hif1a*^*f/f*^/*Nkx2.5*^*Cre/+*^, right), showing compact myocardium (top) and trabeculae (bottom) and quantification of total mitochondria in electron micrographs from E12.5 controls (black bars) and mutants (white bars). Results are expressed as number of mitochondria per tissue area (px^2^). Bars represent mean±SEM (n=4) **B)** RT-qPCR analysis of mitophagy-related genes in E12.5 *Hif1a*-mutant ventricles. Bars (mean±SEM, n=3 for *Bnip3* and *Mxi1* and n=4 for *Nix*) represent fold induction relative to baseline expression in littermate controls (red line). **C)** GSEA enrichment plot for the Hallmark database Oxidative Phosphorylation gene set. The red to blue stripe represents 14406 genes detected as expressed after differential expression analysis, ranked by logFC. Genes at the left side (coloured in red) are more expressed in *Hif1a*/*Nkx2.5* mutants (*Hif1a*^*f/f*^/*Nkx2.5*^*Cre/+*^) and those located at the right side (coloured in blue) are more expressed in control (*Hif1a*^*f/f*^/*Nkx2*^*r+/+*^) littermates. Vertical black lines represent the position of members the Oxidative Phosphorylation gene set along the ranked collection of genes. The green curve represents cumulative enrichment score and its rigth skew indicates that oxidative phosphorylation genes tend to be more expressed in mutants. **D)** Fold change gene expression determined by RNASeq of genes involved in fatty acid uptake and catabolism in *Hif1a*/*Nkx2.5* mutants. Red line represents baseline expression in control littermates. Bars represent mean±SEM (n=2) **E)** Quantification of fatty acid-conjugated carnitines by metabolomic untargeted profiling using MS/MS in control (black bars) and *Hif1a*/*Nkx2.5* (white bars) hearts at E12.5. Bars represent mean±SEM (n=13) In all graphs, bars represent mean±SEM), Student’s t test, \**P*<0.05.

As mature cardiomyocytes rely on FA oxidation (FAO) for ATP production and cardiac performance, one possible metabolic adaptation of *Hif1a* deficient hearts associated with the increased mitochondrial content could be an early utilization of FA to provide sufficient ATP levels in the absence of effective glycolysis. However, genes involved in lipid catabolism did not show altered expression between genotypes at E12.5 in the RNASeq analysis (Fig. 4D). Moreover, carnitine and acyl-carnitine profiling by high performance liquid chromatography and mass spectrometry (HPLC/MS) metabolomics did not present significant alterations in *Hif1a*/*Nkx2.5* mutants (Fig. 4E). This indicates that the decreased glycolysis due to *Hif1a* loss is not associated to a compensatory increase in FAO.

These results demonstrate that reduced HIF1 signaling promotes an increment of cardiac mitochondrial network and suggest the activation of metabolic compensatory mechanisms others than FAO activation upon glycolytic inhibition in *Hif1a* mutant embryos

### Amino acid metabolic program is transiently enhanced in cardiac Hif1α deficient embryos

As indicated above, GO enrichment analysis had identified several metabolic process that could be altered upon deletion of *Hif1a*, some of them related with amino acid metabolism and, specifically, to the “cellular response to amino acid starvation” (Table S2). Complementary functional enrichment analyses allowed to pinpoint more specific functional terms, which suggested alterations in Ala, Leu, Val, Ile, Asn, Asp, Ser, and Gly biosynthesis (Fig. 5A, Table S4). These results lead us to hypothesize about the activation of a metabolic reprogramming towards amino acid oxidative catabolism in embryonic cardiomyocytes, in the absence of effective glycolysis associated to *Hif1a* deletion.

**Figure 5.**
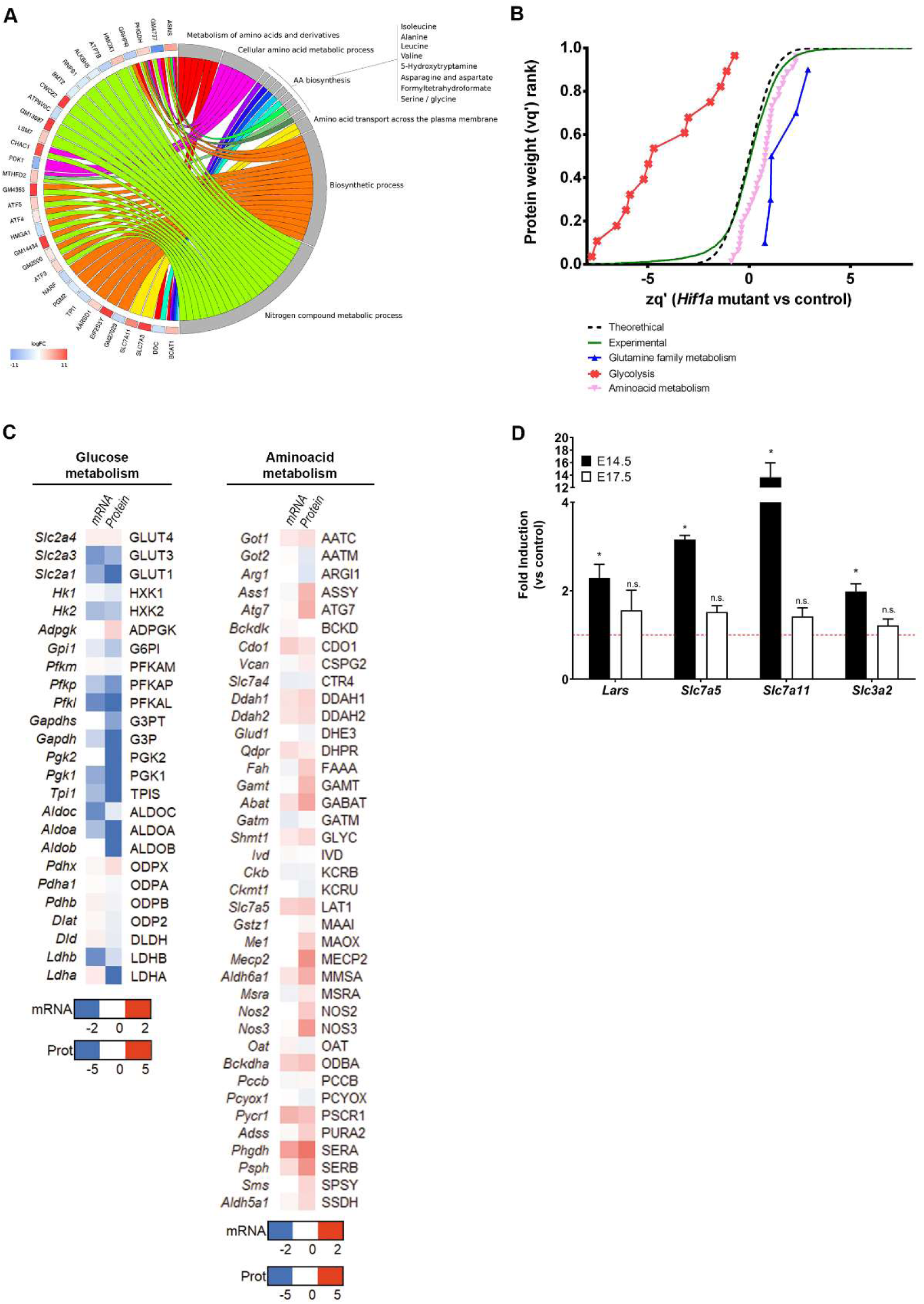
Metabolic adaptations in *Hif1a*-deficient hearts at E12.5. **A)** Circular plot representing logFC values for genes detected as differentially expressed in mutant (*Hif1a*^f/f^/*Nkx2.5*^Cre/+^) embryos, relative to control (*Hif1a*^f/f^/*Nkx2.5*^+/+^), at E12.5, associated to a selection of functional descriptions related to aminoacid metabolism. Functional descriptions were detected with Panther by comparison against the Biological Process component of the Gene Ontology database, and the Panther Pathway and Reactome databases. All functional descriptions were enriched with P value < 0.05. logFC values are color coded: red colour denotes higher expression in mutant samples. **B)** Representation of protein statistical weights (wq’) grouped by functional categories (FDR<1%, n=6) versus protein abundance in *Hif1a*-deficient hearts relative to control embryos (zq’) at E12.5, as determined by MS/MS proteomics. A displacement right from the experimental curve indicates increased pathway in mutant embryos. **C)** Heatmap representation of mRNA (quantified by RNASeq) and Protein (quantified by MS/MS) of components of glucose (left) and amino acid (right) metabolic pathways. Color code indicated in the legend is calculated as the value found in *Hif1a*/*Nkx2.5* mutants relative to control littermates. **D)** RT-qPCR analysis of amino acid transporter gene expression in E14.5 (black bars) and E17.5 (white bars) *Hif1a*-mutant ventricular tissue. Bars (mean±SEM, n=2-4 for E14.5 and n=3 for E17.5) represent fold induction relative to baseline expression in littermate controls (red line). Student’s t test, * pvalue<0.05, ***pvalue<0.005, n.s. non-significant.

To validate this hypothesis, we performed global proteomic analysis in *Hif1a*/*Nkx2.5* embryos and control littermates by E12.5. Quantitative analyses revealed a significant increase in proteins related to amino acid metabolism and, specifically, glutamine family metabolism (Fig. 5B, Table S5). In addition, the observed decrease in proteins related to glycolysis confirmed the inhibited glycolytic gene expression program upon *Hif1a* deletion (Fig. 3). Quantitative results for both gene expression, by RNASeq, and protein abundance, by MS/MS proteomics, therefore correlate to a certain extent, and indicate inhibited glycolysis (Fig. 5C, left) and increased amino acid catabolism (Fig. 5C, right). Specifically, *Hif1a*/*Nkx2.5* mutants showed increased expression and protein levels of genes contributing to anaplerosis of amino acids into Krebs’ Cycle. These contributions (Fig. S5) included aromatic amino acids (Phe, Tyr), polar amino acids (Asn, Asp, Gln, Glu, Ser, and Cys), Pro, Gly and branched-chain amino acids (Val, Leu, and Ile). Urea Cycle was also upregulated, resulting in increased contribution of Arg to Krebs’ Cycle by the generation of fumarate, which can also act as a Krebs’ cycle intermediate. To determine whether this amino acid signature was maintained over time in *Hif1α*-deficient hearts, we analyzed the expression levels of amino acid transporters by RT-qPCR at E14.5 and E17.5 (Fig 5D). The results showed that gene expression levels of several transporters, such as *Lat1* (transporter of Trp, Phe, Tyr and His, but also Met, Val, Leu and Ile (Yanagida et al., 2001)), *Slc7a11* (transporter of Cys (Lim and Donaldson, 2011)), and *Slc3a2* (transporter of Val, Leu and Ile by association with *Slc7a5* (Kanai et al., 1998)), as well as the leucyl-tRNA synthetase *Lars*, were still upregulated by E14.5 in *Hif1a*-deficient hearts, but reached control-like expression levels by E17.5.

This data indicates that upregulation of amino acid transport is transient and suggest that temporary increase of amino acid catabolism and anaplerosis could act as a compensatory mechanism to overcome the loss of glycolytic metabolism upon *Hif1a* loss until the FAO is stablished later in gestation. Moreover, our observations reveal the metabolic flexibility of the embryonic heart to adapt to different substrates for energy supply and support the idea that *Hif1a*-deficient hearts are still able to switch their metabolism towards FAO after E14.5.

### ATF4 signaling is upregulated in *Hif1a*/*Nkx2.5* embryos

The fact that there is a transient upregulation of general amino acid catabolism upon *Hif1a* loss suggests the existence of upstream regulators that are temporally induced in the *Hif1a*/*Nkx2.5* mutant. Amino acid metabolism and its transport is tightly regulated through several pathways, including mTOR (mammalian target of rapamycin), GCN2 (general control non-derepresable 29) and ATF4 (Activating Transcription Factor 4), among others (Bröer and Bröer, 2017). Upstream regulator analysis of our RNASeq data using Ingenuity Pathway Analysis (IPA) in fact shows that ATF4 and CHOP could function as main regulators of a gene set implicated in amino acid metabolism in *Hif1a*-deficient hearts (Fig. 6A and Table S6). ATF4 is a transcriptional regulator that activates the expression of genes involved in amino acid transport and metabolism (Harding et al., 2003) and also respond to nutrient and metabolic stress in hypoxia (Weidemann and Johnson, 2007). *Atf4* gene expression, and mRNA levels of its target genes, such as *Slc7a11*, *Slc7a3*, *Aars*, *Lars* or *Trib3* among others, are positively regulated in our *Hif1a* deletion model by E12.5 (Fig 6B). Interestingly, the transcriptional upregulation of ATF4 was sustained at E14.5, but its expression returned to control hearts levels by E17.5 (Fig. 6C), following a similar expression pattern of amino acid transporters (Fig 5E).

**Figure 6.**
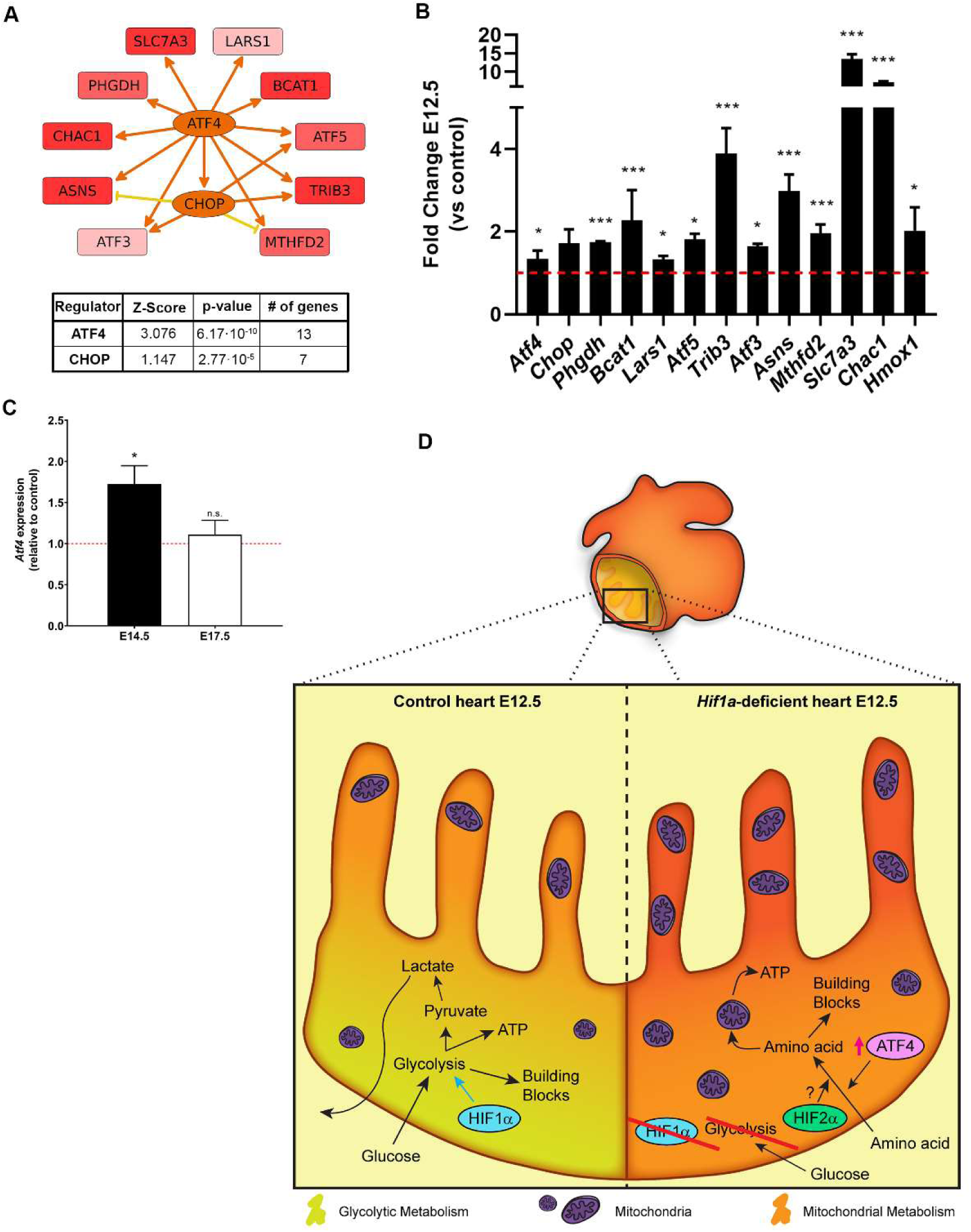
Upstream Regulators of amino acid catabolism activation. **A)** ATF4-centered regulatory network summarizing the interactions between ATF4 itself and CHOP with a collection of genes related to amino acid metabolism, detected as differentially expressed in *Hif1a*-deficient hearts (*Hif1a*^f/f^/*Nkx2.5*^Cre/+^) relative to control hearts (*Hif1a*^f/f^/*Nkx2.5*^+/+^), at E12.5. The graph is a simplified version of a mechanistic network predicted after IPA’s Upstream Regulator Analysis on the complete set of 201 differentially expressed genes. Intensity of red colour in target genes is proportional to logFC. Intensity of orange color in regulator genes (ATF4 and CHOP) is proportional to the predicted activation state. Arow-pointed and flat-headed lines represent positive and negative regulation interactions, respectively. Orange and yellow lines represent congruent and uncongruent connections, respectively, relative to the predicted activation state of regulators. The inset at the right side summarizes z-score value and enrichment P value for ATF4 and CHOP, as well as the number of differentially expressed genes that are regulated by either of them. **B)** Relative expression versus control of genes related to amino acid metabolism downstream of ATF4 determined by RNASeq at E12.5 (n=2). **C)** RT-qPCR analysis *Atf4* gene expression in E14.5 (black bar) and E17.5 (white bar) *Hif1a*-mutant ventricular tissue. Bars (mean±SEM, n=5 for E14.5 and n=3 for E17.5) represent fold induction relative to baseline expression in littermate controls (red line). Student’s t test, * pvalue<0.05, ***pvalue<0.005, n.s. non-significant.**D)** Model representing the embryonic myocardium by E12.5. Compact myocardium is mainly glycolytic (yellowish), by the action of HIF1 signaling, while trabeculae rely more on mitochondrial metabolism (orange) in control embryos (left). In *Hif1a*-mutants (right), the whole myocardium relies on mitochondrial metabolism and present higher mitochondrial content, using amino acids as energy source through the activation of ATF4 signaling.

Taken together, these data suggest that dynamic ATF4 pathway activation could be regulating, the transient increase in the amino acid catabolic program shown in *Hif1*-deficient embryos, hence contributing to the metabolic adaptation in the absence of glycolysis during cardiac development.

In summary, our data demonstrate that HIF1 signaling in NKX2.5 cardiac progenitors is dispensable for proper heart formation and that the absence of *Hif1a* triggers a cardiac metabolic reprogramming, enhancing temporal amino acid catabolism to ensure sufficient ATP and biosynthetic precursors to sustain cardiac growth and function even in the absence of glycolysis (Fig. 6D). Importantly these adaptations might be relevant in the adulthood under pathological scenarios associated with oxygen signaling like pulmonary hypertension or cardiomyopathy towards the development of new drugs against new metabolic targets.

## DISCUSSION

Here, we describe that *Hif1a* loss in NKX2.5 cardiovascular progenitors causes glycolytic program inhibition in the compact myocardium (CM) by E12.5, without compromising normal cardiac development and embryonic viability. Our results show that upon *Hif1a* deletion, the embryonic myocardium conserves FAO capacity but exhibits the ability to activate metabolic programs oriented to amino acid catabolism, together with an increase in the mitochondrial content by E12.5. Taken together, our findings point out the metabolic versatility of the embryonic heart and conciliate the discrepancies from previous deletion models of *Hif1a* in cardiovascular progenitors.

### Integration with previous *Hif1a* deletion model in the embryonic heart

As outlined in the introduction, there is a lack of consensus between previous reports on cardiac embryonic mouse models of *Hif1a* loss. On the one hand, the use of Mlc2vCre, a less robust and not homogenously expressed driver within the myocardium as *Nkx2.5*, might lead to heterogeneous recombination pattern and hence not complete abrogation of HIF1 signaling in cardiomyocytes (Li et al., 2011) that could cause the adult phenotype described by Huang and collaborators (Huang et al., 2004). Thus, the use of different Cre drivers and its differences in terms of recombination pattern and efficiency keeps off any kind of comparison between genetic models. On the other hand, previous reports from Perriard and Zambon/Evans’ group (Guimarães-Camboa et al., 2015; Krishnan et al., 2008), despite of the Cre driver applied, use a null allele of *Hif1a* in combination with a floxed allele. The *Hif1a* haploinsufficiency of these models outside heart NKX2.5 territories might cause extracardiac affections, such as vascular or placental, that could significantly influence the described phenotype. Indeed, the use of a *Hif1a*-null allele has been reported to cause cardiovascular malformations associated with maternal diabetes (Bohuslavova et al., 2013). In this regard, the exhaustive characterization by Guimaraes-Camboa et al. at gene expression level extensively overlaps with our RNASeq data, except in terms of stress and apoptotic pathways, upregulated only in the null-allele context.

Moreover, we have also analyzed a *Hif1a* null/floxed model in NKX2.5 progenitors in parallel with the double floxed model described here. We found that although the glycolytic inhibition by E14.5 is comparable in both models (data not shown), only the null/floxed mice exhibited embryonic lethality (5% retrieved versus 25% expected, pvalue 0.0029, n=7 litters), in contrast with viability of the double floxed mice. This observation, together with our results, support the notion that *Hif1a* is dispensable for cardiac development, while an extensive comparison, in terms of gene expression in placental and vascular embryonic tissue, between the double floxed and the null/floxed models would be necessary to exclude extracardiac influences of *Hif1a* deficiency impacting on heart development as reported by Guimaraes-Camboa and colleagues.

### Amino acid catabolism and metabolic versatility of the embryonic heart

A key finding of our investigation is the fact that the embryonic myocardium is able to upregulate alternative metabolic pathways (amino acids catabolism), and to promote mitochondrial enrichment that could support the ATP demand upon glycolytic inhibition subsequent to *Hif1a* loss. The use of amino acids as cardiac metabolic fuel has been proposed mainly in oxygen-deprived scenarios (Bing et al., 1954; Julia et al., 1990). Amino acids provide, by deamination, carbon skeletons that can be converted into pyruvate, alpha-ketoglutarate, succinyl-CoA, fumarate, oxalacetate, Acetyl-CoA and Acetoacetyl-CoA, all of them metabolites that can be incorporated into the Krebs Cycle (Evans and Heather, 2016; Neubauer, 2007). As detailed above, our *Hif1a*-defficient model upregulates, both at the transcriptional and protein level, a variety of amino acid transporters, biosynthetic and catabolic enzymes, that can replenish the Krebs Cycle’s upon glucose deprivation. Moreover, the fact that this upregulation is accompanied by an increase in mitochondrial content indicates that the embryonic heart, in the absence of *Hif1a*, readapts its metabolism to maintain enough ATP levels and building blocks, without compromising the normal protein synthesis required for myocardium development and embryo viability.

Interestingly, this adaptation is transient and reversible, as seen by the control-like levels of amino acid transporters transcripts found at later stages by E17.5, without precluding the embryonic metabolic switch towards FAO previously described (Menendez-Montes et al., 2016). This shows that the embryonic myocardium has the plasticity to modulate its metabolism to adapt to the energetic demand and nutrient availability. Moreover, in addition to this interesting role of amino acid catabolism activation in the embryonic context, the use of amino acids as an alternative energy source could be an interesting option to achieve cardioprotection and recovery after cardiac injury. In this regard, some of the enzymes upregulated in our massive screenings in *Hif1a*-deficient hearts are involved in Ser biosynthesis and one-carbon cycle, including *Phgdh*, *Psph* and *Shmt1*. These pathways have been previously described to increase glutathione levels and protect the heart against oxidative stress (Zhou et al., 2017), also in a context of myocardial hypertrophy (Padrón-Barthe et al., 2018).

### Origin of catabolized amino acids in *Hif1a*-deficient hearts

An interesting open aspect of the metabolic adaptation exhibited by the *Hif1a*-deficient hearts is the source of amino acid supply during glycolytic inhibition upon *Hif1a* loss. In this regard, two potential sources could be considered. First, protein-forming amino acids could be recycled through autophagy. This hypothesis is reasonable considering the context of the embryonic heart, where protein turnover, specially transcription factors, happens fast and at a high rate (Merz et al., 1981). Moreover, a positive nitrogen balance has been reported in both adult rat and human hearts, indicative of rapid turnover of tissue proteins (Sprinson and Rittenberg, 1949). Interestingly, our *Hif1a* mutant embryos showed increased transcription of *p62*, a cargo-recognizing protein involved in autophagic degradation of cellular proteins (Lim et al., 2015). While an extensive characterization analyzing autophagy, pro and anti-autophagic signaling pathways and protein labelling and turnover would be needed to further investigate this hypothesis, the fact that autophagy could be involved in this metabolic adaptation suggest an exciting link between cardiac metabolism, hypoxia and autophagy.

Another possible source of amino acids in *Hif1a*-deficient hearts is fetal circulation. Even though an extensive characterization of fetal blood nutrient content over gestation has not been reported, the transcriptional increase in several membrane amino acid transporters observed in our mutants suggest that *Hif1a*-deficient cardiomyocytes could be obtaining them directly from embryonic circulation. Interestingly, cardiac amino acids uptake in human subjects infused intravenously with protein hydrolysate increases by 245% (Bing et al., 1954), showing that the heart can respond to blood amino acids levels. Moreover, the regulation of amino acid transporters expression in the placenta is essential for maintaining high levels of amino acids in the fetal blood to sustain embryo growth (Díaz et al., 2014). In this regard, an increased cardiac uptake of amino acids in the *Hif1a*-deficient embryo could result in increased amino acids supply through the placenta that might response to some secreted cues in the absence of cardiac HIF1 signaling.

### Molecular determinants of amino acid catabolism activation

Amino acids metabolism and transport is tightly regulated through several pathways, including mTOR, GCN2 and G-protein-coupled receptors, among others (Bröer and Bröer, 2017). ATF4 is a transcriptional regulator that activates the expression of genes involved in amino acid transport and metabolism (Harding et al., 2003) and also respond to nutrient and metabolic stress in hypoxia (Weidemann and Johnson, 2007). Atf4 gene expression is positively regulated in our *Hif1a* deletion model at both the transcriptional (Fig. 6C) and at the protein level (Fig. S6A). In addition, upstream regulators analysis identified ATF4 and CHOP (Ddit3) as putative regulators of amino acid metabolism in *Hif1a*-deficient hearts (Fig. 6A). ATF4 has been identified as an essential factor for amino acid starvation, by activating gene expression of genes containing amino acid response elements (AARE) (Zhang et al., 2010) and regulates CHOP expression (Averous et al., 2004).

Loss of HIF1α signaling results in downregulation of glycolytic enzymes expression (Fig. 2) and glycolytic inhibition and glucose deprivation has been shown to cause activation of Unfolded Protein Response (UPR) (Badiola et al., 2011; Ikesugi et al., 2006; Vavilis et al., 2016). In our deletion model, reduced glycolysis was accompanied by increased in UPR activation (Fig. S6B), suggesting that *Hif1a* deletion could contribute to ATF4 activation through glycolytic inhibition and UPR activation.

Interestingly, the *Hif1a* deletion model by Guimaraes-Camboa and collaborators (Guimaraes-Camboa et al., 2015) also shows increased ATF4 signaling in *Hif1a*-deficient hearts, supporting ATF4 as one of the main regulators of the described metabolic adaptation upon loss of effective glycolysis in our *Hif1a*-deficient hearts. Moreover, since ATF4 is upregulated in both animal models, and considering the lack of lethality of our floxed/floxed mice, the upregulation of ATF4 stress pathway does not seem to be responsible of the embryonic lethality reported by Zambon’s/Evan’s groups.

Additionally, HIF1 signaling has been also described to directly interact with MYC pathway in cancer settings (Gordan et al., 2007b). Specifically, HIF1 disrupts the formation of MYC/MAX complexes by increasing Mxi1 expression. Therefore, in a context of downregulated HIF1 signaling, MYC-mediated transcription could be activated. Moreover, cMYC promotes amino acid metabolism, including the expression of Slc7a5 (*Yue et al., 2017*), involved in the transport of large neutral amino acids, such as Arg, Leu or Tyr. Interestingly, *Mxi1* transcription is significantly reduced in our mutant mice (Fig 4B), supporting the idea that HIF-mediated repression of MYC/MAX complexes could be abrogated in the *Hif1a*-deficient model and hence MYC could be an additional potent contributor of amino acid catabolic pathways. This observation sets the basis for further research in metabolic adaptations by MYC in the absence of active HIF1 signaling.

### Possible role of HIF2a and future directions

Finally, in the absence of active HIF1 cascade, HIF2alpha, an alternative HIF alpha isoform able to form functional heterodimers with ARNT could play a compensatory role. To explore this possibility, we analyzed HIF2α abundance by western blot finding a protein induction by E12.5 in the *Hif1*/*Nkx2.5*-deficient hearts and comparable levels at E14.5 between control and mutants. A similar induced expression was observed for HIF2 target gene, PAI1 (Fig S7). Interestingly, glucose deprivation has been shown to induce HIF2 signaling in an acetylation-dependent manner (Chen et al., 2015). We hypothesize that this adaptive induction of HIF2 signaling could be related with the transcriptional activation of amino acid catabolic program observed in the cardiac *Hif1a*-defficient model. On one hand HIF2α has been described to activate cMYC in cancer cells (Gordan et al., 2007a) and MYC could modulate amino acid catabolism as discussed above. On the other hand, HIF2α has been also involved in the direct expression of *Slc7a5* by binding to the proximal promoter of the gene in renal clear cell carcinoma WT8 cells, as well as in lung and liver cells (Corbet et al., 2014; Elorza et al., 2012; Zhang et al., 2020). Additionally, HIF2 signaling has been related to the expression of other amino acid metabolism-related genes, such as *Mthfd2* or *Atf3* (Green et al., 2019; Turchi et al., 2008). Hence, these preliminary results suggest that HIF2 could be an additional potent inductor of cardiac amino acid catabolic pathways, opening new research horizons in our lab to study the role of HIF2 as an important player modulating cardiac metabolic compensatory responses in the absence of effective HIF1 signaling.

## MATERIALS AND METHODS

### Animal care and housing

*Hif1a*^*flox/flox*^ (Ryan et al., 2000)mice were maintained on the C57BL/6 background and crossed with mice carrying *Nkx2.5Cre* recombinase (Stanley et al., 2002) or *TnTCre* recombinase (Jiao et al., 2003) in heterozygosity. *Hif1a*^*flox/flox*^ homozygous females were crossed with double heterozygous males and checked for plug formation. Mice were housed in SPF conditions at the CNIC Animal Facility. Welfare of animals used for experimental and other scientific purposes conformed to EU Directive 2010/63EU and Recommendation 2007/526/EC, enforced in Spanish law under Real Decreto 53/2013. Experiments with mice and embryos were approved by the CNIC Animal Experimentation Ethics Committee.

### Genotyping

Genotyping was performed as previously described (Menendez-Montes et al., 2016) using the following primers (Sigma Aldrich; USA) for *Hif1a floxed* alleles: 5’ CGTGTGAGAAAACTTCTGGATG 3’ and 5’ AAAAGTATTGTGTTGGGGCAGT 3’. For *Hif1a* null allele detections the primers were: 5’ GCCCATGGTAAGAGAGTAGGTGGG 3’ and 5’ 5’ AAAAGTATTGTGTTGGGGCAGT 3’. For Cre alleles genotyping, the following primers were used: Nkx2.5: 5’ GCCCTGTCCCTCAGATTTCACACC 3’, 5’ GCGCACTCACTTTAATGGGAAGAG 3’ and 5’ GATGACTCTGGTCAGAGATACCTG 3’ and *cTnT*: 5’ TACTCAAGAACTACGGGCTGC 3’ and 5’ GCACTCCAGCTTGGTTCCCGA 3’.

### Embryo extraction

Embryos were extracted after pregnant female euthanasia by CO_2_ inhalation and head and liver were removed. Embryos were then fixed overnight at 4ºC in 4% PFA solution (RT15710, Electron Microscopy Sciences; USA). After fixation, embryos were dehydrated, embedded in paraffin and sectioned at 5µm for immunostaining and histological purposes and at 10µm for *in situ* hybridization. For mitochondrial activity staining, embryos were directly embedded in OCT and snap frozen in liquid nitrogen and sectioned at 10µm using a cryostat.

### Histological and immunohistochemical analysis

Histological sample processing and immunostaining was performed as previously described (Menendez-Montes et al., 2016). The primary antibodies used in this study were: HIF1α (NB100-479, Novus Biologicals; USA and GTX30647, Genetex, USA); cTnT (CT3, Developmental Studies Hybridoma Bank; USA); BrdU (347580, BD Biosciences; USA); Cy3-conjugated Smooth Muscle Actin (C6198, Sigma Aldrich; USA) and GLUT1 (Cat. No. 07-1401, Millipore, USA). FITC-WGA (W32466, Life Technologies; USA) was used to stain cell membrane. Nuclei were stained with DAPI (Millipore; USA).

### Quantification of histological and immunostained sections

HE staining was quantified as described previously (Menendez-Montes et al., 2016) using ImageJ (Rasband, 2015). For fluorescence intensity analysis in cardiomyocyte nuclei, our own pipeline for CellProfiler software was employed (Lamprecht et al., 2007). Briefly, cell nuclei were segmented and subsequently filtered by cTnT positive cytoplasmic staining. After filtration, HIF1α channel intensity was measured. For WGA-based area quantification, 25 cardiomyocytes in RV, LV and IVS were manually quantified. Only cardiomyocytes in cross-sectional area at the level of the nucleus were quantified.

### RNA extraction, cDNA synthesis and RT-qPCR

RNA extraction from embryonic hearts, cDNA synthesis and quantitative PCR were performed as previously described (Menendez-Montes et al., 2016). Primers are available under request.

### Probe synthesis and in situ hybridization

General probe synthesis, purification and in situ hybridization steps were followed according with our previous protocol (Menendez-Montes et al., 2016). For the probe synthesis, the following primers were used: *Glut1* 5’ GGACTTTGATGGCTCCAGAA 3’ and 5’ GAGTGTCCGTGTCTTCAGCA 3’, *Pdk1* 5’ CTGGGTTTGGTTACGGATTG 3’ and 5’ GCCAGCTACTCCACGTTCTT 3’ and *Ldha* 5’ GGAAGGAGGTTCACAAGCAG 3’ and 5’ CTGCAGTTGGCAGTGTGTCT 3’.

### Electron microscopy and micrograph quantification

Embryonic hearts were processed for transmission electron microscopy following the standard procedures. Briefly, after overnight fixation in 3% glutaraldehyde/4%PFA, samples were refixed in 1% osmium tetroxide and embedded in epoxy resin. 60nm sections were counterstained with uranyl acetate and lead citrate and imaged using a JEOL JEM1010 (100 KV) transmission electron microscope. Control and mutant embryonic hearts from three independent litters were analyzed. For quantification of mitochondria and lipid droplets, ten images of compact myocardium and ten of trabeculae were taken at 5000x magnification. Mitochondria and droplets were counted manually by blinded observers using the ImageJ CellCounter plugin. Values were normalized to the total tissue area, in pixels, excluding extracellular areas in the image.

### Protein extraction and Western Blot

Embryonic hearts were homogenyzed using RIPA buffer and a TissueLyser in presence of protease and phosphatase inhibitors (Inhibitor cocktail (11697498001, Roche, Switzerland) and 1μM sodium ortovanadate). After clarification by centrifugation, protein concentration was measured using Pierce BA Protein Assay kit (23227, Thermo Scientific; USA) following manufacturer instructions. 30μg of protein were denaturized at 95ºC for 5 min, loaded on an 8% polyacrylamide SDS-PAGE gel and run at 120V for 90min. Subsequently, samples were transferred to a nitrocellulose membrane by wet transfer at 400mA for 2h. Membranes were blocked with 5% BSA for 1h and incubated with primary antibodies O/N at 4ºC. Next day, membranes were washed in TBS-T buffer and incubated with the corresponding HRP-conjugated secondary antibodies (Dako, Denmark) at 1:5000 dilution for 1h at RT. After washing, signal was developed using ECL Primer Western Blotting Detection Reagent (Amersham; UK) and detected by a LAS-3000 imaging system (Fujifilm; USA). The primary antibodies used in this study were: anti-HIF2α[ep190b] (NB100-132, Novus Biologicals) dilution 1:200, anti-HIF1α (10006421, Cayman; USA) dilution 1:200, anti-PAI-1 (sc-5297, Santa Cruz) dilution1:500, anti-ATF4 (11815, Cell Signalling; USA) dilution 1:500, anti-vinculin (V4505, Sigma-Aldrich; USA) dilution1:5000, anti-αtubulin[DM1A] (ab7291,Abcam; UK) dilution 1:1000 and anti-SMA cy3 (C6198, Sigma-Aldrich; USA) dilution 1:1000.

### RNA-Seq and bioinformatics analysis of gene expression

RNA-Seq data processing and differential expression analyses (n=2 per group) were performed as previously described (Menendez-Montes et al., 2016). For differential expression analysis, only genes expressed with at least at 1 count per million in at least in 2 samples were considered. Changes in gene expression were considered significant if associated with Benjamini-Hochberg adjusted P value < 0.055. Functional enrichment analyses were performed with Gorilla (Eden et al., 2009), Panther (Thomas et al., 2003), IPA (Qiagen, USA), REVIGO (Supek et al., 2011) and GSEA (Subramanian et al., 2005). Enriched functional terms were filtered by applying P value thresholds described in the corresponding Table or Figure captions. Circular plots summarizing logFC values for genes, and their association to enriched functional terms were generated with GO plot (Walter et al., 2015).

### Metabolomics untargeted profiling analysis by HPLC

Metabolites from single E12.5 embryonic hearts were extracted by polypropylene pestle homogenization using 0.1% ammonium acetate:methanol (1:1 v/v) solution containing 5mM BHT. Proteins were precipitated by cold 1:1 methanol-ethanol and supernatant was dried in speedvac (Savant SPD131DDA, ThermoFisher; USA) for 2h at RT. Metabolomics analysis was performed using an Ultimate 3000 HPLC equipped with a Merck SeQuant ZIC-HILIC column (150 × 1 mm, 3.5µ) thermostated at 45°C and coupled to a LTQ Orbitrap XL MS (ThermoFisher; USA). Metabolites were eluted at 180µL/min with A: water with 0.1% formic acid (FA) and B: acetonitrile with 0.1% FA. Data were collected in positive and negative ESI ion full scan mode and CID-DDA mode, from 70 to 1000 m/z, at 60000 resolution. Data processing was carried-out using Compound Discoverer (ThermoFisher; USA) with the Metaboprofiler node (Röst et al., 2016). Putative identification was performed using Ceu Mass Mediator (Gil-De-La-Fuente et al., 2019).

### Proteomics analysis

Proteins from pellets after metabolite extraction were pooled in groups of four, treated with 50mM iodoacetamide (IAM) and digested with trypsin using the Filter Aided Sample Preparation (FASP) digestion kit (Expedeon) (Wiśniewski et al., 2011) according to manufacturer’s instructions. Dried peptides were labeled with iTRAQ-8plex according to manufacturer’s instructions, desalted on OASIS HLB extraction cartidges (Waters Corp.), separated into 4 fractions using the high pH reversed-phase peptide fractionation kit (Thermo) and dried-down before MS analysis on an Orbitrap Fusion Tribrid mass spectrometer (Thermo Fisher Scientific, Bremen, Germany) (García-Marqués et al., 2016a). Peptide identification, quantification and systems biology analysis was performed as in (García-Marqués et al., 2016) Significant abundance changes of proteins or homogeneous categories of KO mice compared to controls were detected at 1% FDR.

### Adult mice echocardiography and analysis

5 months-old mice were anesthetized using 1.5% isoflurane at a flow rate of 1L/min. Once anesthetic plane was reached, cardiac images were acquired using a MS400 probe, at 30MHz for 2D and M mode images and 24MHz for Color and Pulsed Doppler modes, using an ultrasound scanner VEVO2100 (Visualsonics, Canada).

### Statistical analysis and data representation

For histological, immunohistochemical quantifications, electron microscopy and RT-qPCR, values were pooled for embryos with the same genotype from independent litters and analyzed by the indicated statistical test using SPSS software (IBM; USA), with statistical significance assigned at P ≤0.05. Values were represented as mean±SEM using GraphPad Prism (GraphPad; USA).

### Data access

RNASeq data: The accession number for the RNAseq data reported in this paper is GEO:

Proteomics data: The data set (raw files, protein databases, search parameters and results) is available in the PeptideAtlas repository (PASS01235).

## ACKNOWLEDGMENTS

Authors thank Lorena Flores for echocardiography technical assistance, Raquel Baeza and Mercedes de la Cueva for animal housing and handling and to CNIC Microscopy, Genomics and Histology Core Facilities for technical assistance. We thank Miguel Torres Sanchez, Jose Antonio Enriquez, Luke Szweda and Hesham Sadek for critical discussion of the manuscript.

## SOURCES OF FUNDING

This project has been supported by Fundación Centro Nacional de Investigaciones Cardiovasculares Carlos III (CNIC) and by grants to SM-P from the European Research Council: FP7-PEOPLE-2010-RG_276891, Fundación TV3 La Marató: 201507.30.31, Comunidad de Madrid (CAM) and European Union (EU): B2017/BMD-3875; Instituto de Salud Carlos III: PI17/01817, Universidad Francisco de Vitoria (UFV) and LeDucq Foundation. IMM was supported by La Caixa-CNIC and Fundacion Alfonso Martín Escudero fellowships, TAG was supported by a predoctoral award granted by CAM/EU and UFV: PEJD-2018-PRE/SAL-9529 and SM-P by a Contrato de Investigadores Miguel Servet (CPII16/00050) and UFV.

## DISCLOSURES

None

## Notes

### Competing Interest Statement

The authors have declared no competing interest.

