## Supplemental legends and figures for "Activation of amino acid metabolic program in response to impaired glycolysis in cardiac HIF1 deficient mice"

**SUPPLEMENTARY MATERIAL**

**Supplementary table legends**

**Table S1. RNA-Seq analysis of gene expression in E12.5 *Nkx2.5/Hif1a* deficient hearts.** Total detected transcript quantification and differentially expressed genes (DEG) in control (*Hif1a*$^{f/f}$/*Nkx2.5*$^{+/+}$) and *Hif1a/Nkx2.5* mutant (*Hif1a*$^{f/f}$/*Nkx2.5*$^{Cre/+}$) hearts at E12.5. The table is composed of two spreadsheets that describe raw and normalized expression values for each gene in each sample, averaged values per condition and for all samples, as well as fold change, logFC, P value and Benjamini-Hochberg adjusted P value, in a mutant versus control comparison. The first spreadsheet describes expression values for 201 genes detected as differentially expressed with Benjamini-Hochberg adjusted P values < 0.055. 83 genes were upregulated (red background) in *Hif1a*-deficient hearts versus control littermates, and 118 genes were downregulated (blue background). The second spreadsheet describes expression values for the whole collection of 14406 genes detected in the RNA-Seq experiment.

**Table S2. Gorilla/REVIGO Gene Ontology Analysis of differentially expressed genes.** Over-represented Biological Process GO terms associated to the collection of 201 differentially expressed genes (adjusted P value < 0.055), as detected with GOrilla, with p_value < 0.001 (default threshold). The table describes, for each GO term, the number of mapped annotated genes in the reference data set, the number of mapped annotated genes in the target set, its fold enrichment and P value. Dispensable terms were eliminated after processing the list with REVIGO.

**Table S3. Gene Set Enrichment Analysis.** Enriched gene sets from the Hallmark, Biocarta and Reactome databases associated to the collection of 14,406 expressed genes, as detected with GSEA, with P value < 0.05. The table describes, for each gene set, the calculated normalized enrichment score and associated P value.

**Table S4. Functional enrichment analysis with PANTHER.** Over-represented Reactome pathways, PANTHER pathways and Biological Process Gene Ontology terms associated to the collection of 201 differentially expressed genes (adjusted P value < 0.055), as detected with PANTHER, with p_value < 0.05. The table describes, for each pathway or GO term, the number of mapped annotated genes in the reference data set, the number of mapped annotated genes in the target set, its fold enrichment and associated P value.

**Table S5. Proteomics analysis of control and *Hif1a*-deficient embryonic hearts at E12.5.** The table shows the list of proteins quantified by mass spectrometry analysis. ZQ (KO/WT) values are standardized log2-ratio averages of proteins from mutant (*Hif1a*$^{f/f}$/*Nkx2.5*$^{Cre/+}$) samples relative to embryos (*Hif1a*$^{f/f}$/*Nkx2.5*$^{+/+}$), merged from the 3 biological replicates. 1% FDR is used as criterion for protein statistically significant abundance change.

**Table S6. Ingenuity Pathway Analysis (IPA).** The table presents, in four spreadsheets, functional enrichment results generated with IPA for the set of 201 genes detected as differentially expressed (adjusted p-value < 0.055), for the following analysis types: Canonical Pathways, Upstream Regulators, Diseases and Biofunctions (Downstream Effect analysis) and Regulator Effects. The key parameters provided for the first three analysis types describe enrichment significance (p-value and Benjamini-Hochberg adjusted p-value) and activation z-score (a prediction of the activity of the pathway, regulator or function; in this case, positive and negative values denote higher activity in mutant or control samples, respectively). Absolute z-score values higher than 2 are considered relevant. The fourth analysis type, Regulator Effects, aim at integrating Upstream and Downstream Effect analysis by constructing networks that combine regulators, target genes and their associated functions.

### Supplementary figures

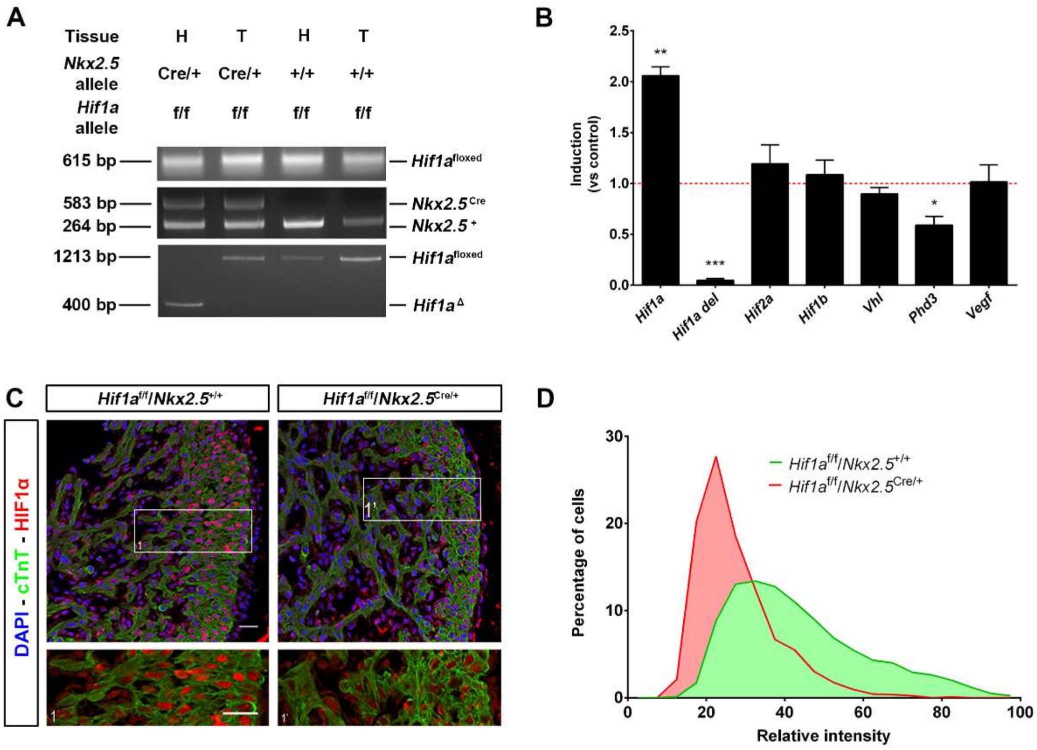


**Figure S1. Deletion efficiency of *Hif1a/Nkx2.5* mutants. A)** Agarose electrophoresis showing PCR products from heart (H) and tail (T) tissue of E12.5 mutant embryos (*Hif1a*$^{f/f}$*/Nkx2*$^{rCre/+}$, lanes 1and 2) and controls (*Hif1a*$^{f/f}$*/Nkx2.5*$^{+/+}$, lanes 3 and 4). Top gel: floxed (615 bp) allele of the *Hif1a* gene. Middle gel: wild-type (264 pb) and Cre (583 bp) alleles of the Nkx2.5 gene. Bottom gel: processed *Hif1a* allele after Cre-mediated recombination (400bp) and unprocessed allele (1213bp). **B)** RT-qPCR quantification of *Hif1a,* exon 2 (floxed) from *Hif1a*, *Hif2a*, *Hif1b¸ Vhl, Phd3 and Vegf* transcripts in E14.5 *Hif1a*- mutant hearts. Bars (mean±SEM, n=3-6) represent fold induction relative to baseline expression in littermate controls (red line). *pvalue<0.05; **0.01<pvalue<0.05; ***pvalue<0.005, Student's t test. **C)** HIF1α immunofluorescence at E12.5 in control and mutant embryos (Dapi staining shows nuclei in blue, Troponin T in green and HIF1α in red). Scale bars, 20µm. **D)** Representative analysis of cardiomyocyte HIF1α nuclear protein expression intensity, quantified by immunohistochemical staining of heart sections from an E12.5 control embryo (green curve) and a *Hif1a*-null littermate (red curve).

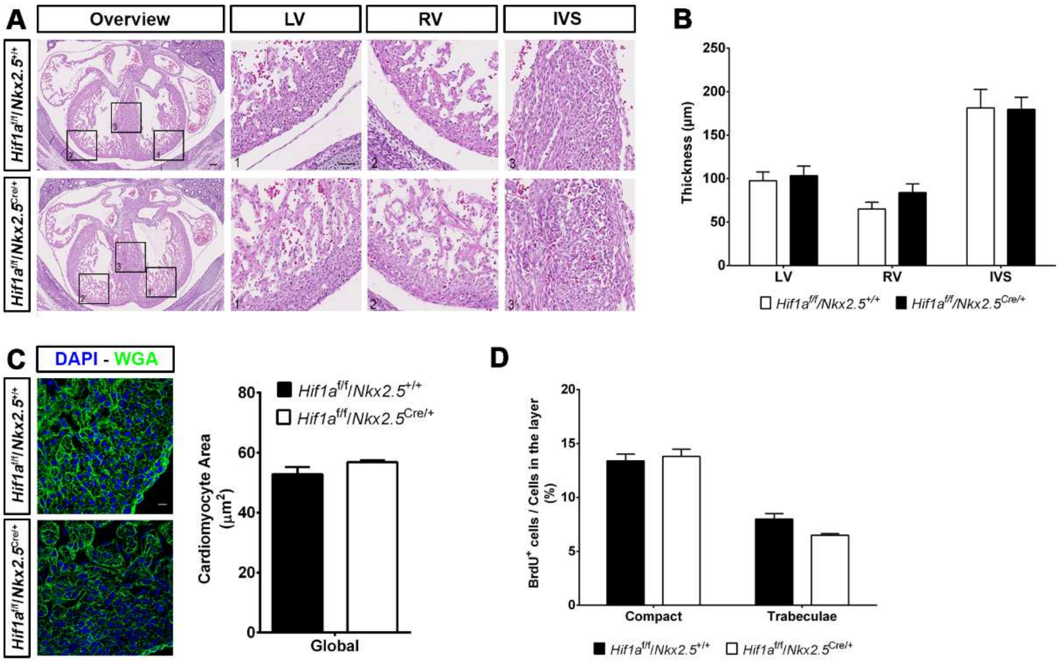


**Figure S2. Embryonic analysis of *Hif1a/Nkx2.5* mutants at E14.5. A)** E14.5 control (*Hif1a*$^{f/f}$*/Nkx2*$^{r+/+}$, up) and mutant (*Hif1a*$^{f/f}$*/Nkx2.5*$^{Cre/+}$, down) embryos stained with hematoxylin and eosin (HE). Scale bars represent 100µm (overview) and 20µm (insets). **B)** HE quantification of ventricular walls and interventricular septum width in E14.5 control (black bars, n=8) and mutant (white bars, n=8) embryos. **C)** LV magnifications of E14.5 control (up and black bar, n=3) and *Hif1a*-deficient (down and white bar, n=3) heart sections stained with wheat germ agglutinin (WGA) and quantification of cardiomyocyte cross-sectional area. **D)** Quantification of BrdU immunostaining, represented as percentage of BrdU$^{+}$ cells in the compact myocardium and trabeculae of E14.5 control (black, n=3) and *Hif1a/Nkx2.5* mutant (white, n=3) embryos. In all graphs, bars represent mean±SEM, Student's t test.

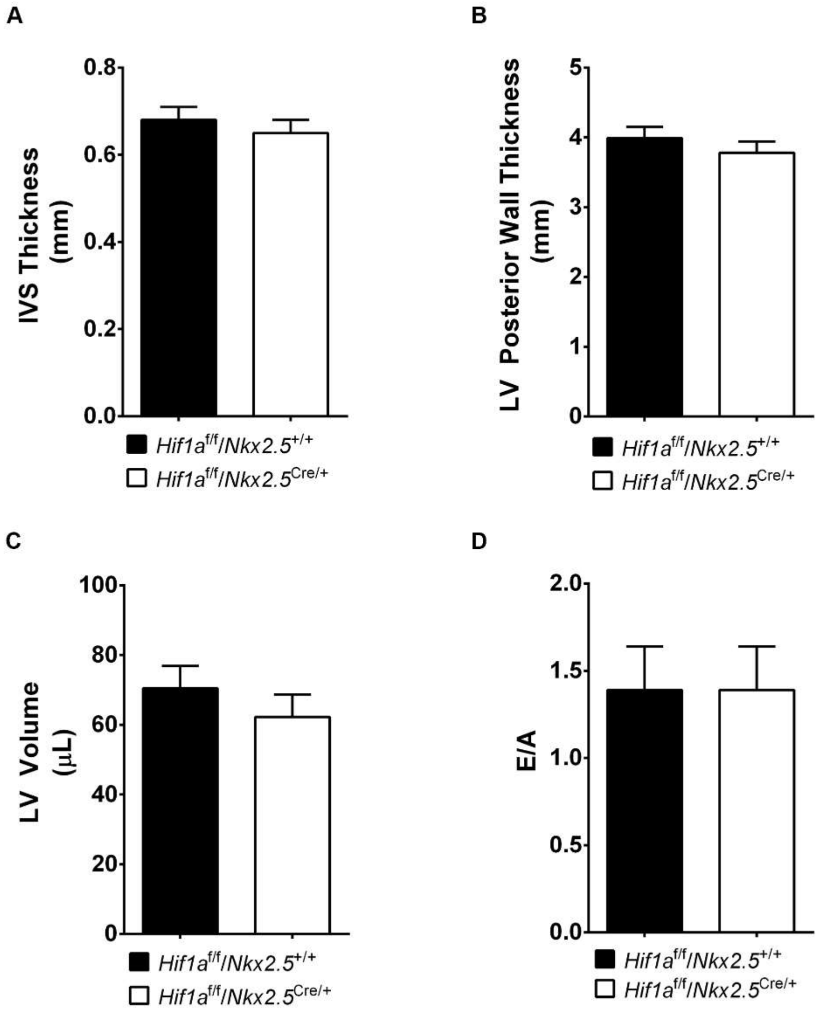


**Figure S3. Morphological and functional echocardiographic parameters in *Hif1a/Nkx2.5* adult mutants.** Quantification of IVS thickness **(A)**, LV posterior wall thickness **(B)**, LV volume **(C)** and diastolic function E/A **(D)** in control ($Hif1a^{f/f}/Nkx2^{r+/+}$, black bars n=8-9) and *Hif1a/Nkx2.5* ($Hif1a^{f/f}/Nkx2.5^{Cre/+}$, white bars n=10-11) at 5 months of age. Bars represent mean±SEM, Student's t test. IVS: interventricular septum; LV: left ventricle; E/A: E wave/A wave.

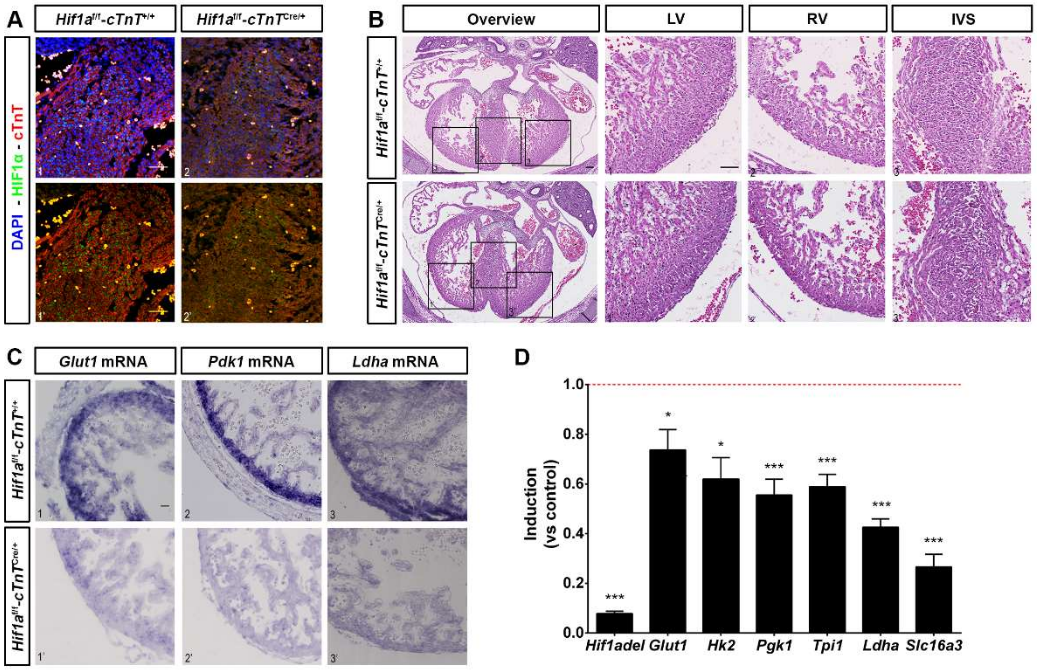


**Figure S4. Analysis of *Hif1a/cTnT* embryos at E14.5. A)** IVS magnifications of E14.5 cardiac sections in control (*Hif1a*$^{f/f}$/*cTnT*$^{+/+}$) and *Hif1a/cTnT* mutant (*Hif1a*$^{f/f}$/*cTnT*$^{Cre/+}$) embryos stained for HIF1α immunofluorescence (Dapi staining shows nuclei in blue, Troponin T in red and HIF1α in green). Scale bars, 20µm. **B)** E14.5 control and *Hif1a/cTnT* mutant embryos stained with hematoxylin and eosin (HE). Scale bars represent 100µm (overview) and 20µm (insets). **C)** RV magnifications of E14 control and *Hif1a/cTnT* embryos analyzed by in situ hybridization against *Glut1* (left), *Pdk1* (middle) and *Ldha* (right) mRNA expression. Scale bar represent 20µm. **D)** RT-qPCR analysis of glycolytic genes in E14.5 *Hif1a/cTnT* mutant ventricles. Bars (mean±SEM, n=3) represent fold induction relative to baseline expression in littermate controls (red line). Student's t test, *pvalue<0.05; ***pvalue<0.005. RV: right ventricle; LV: left ventricle; IVS: interventricular septum.

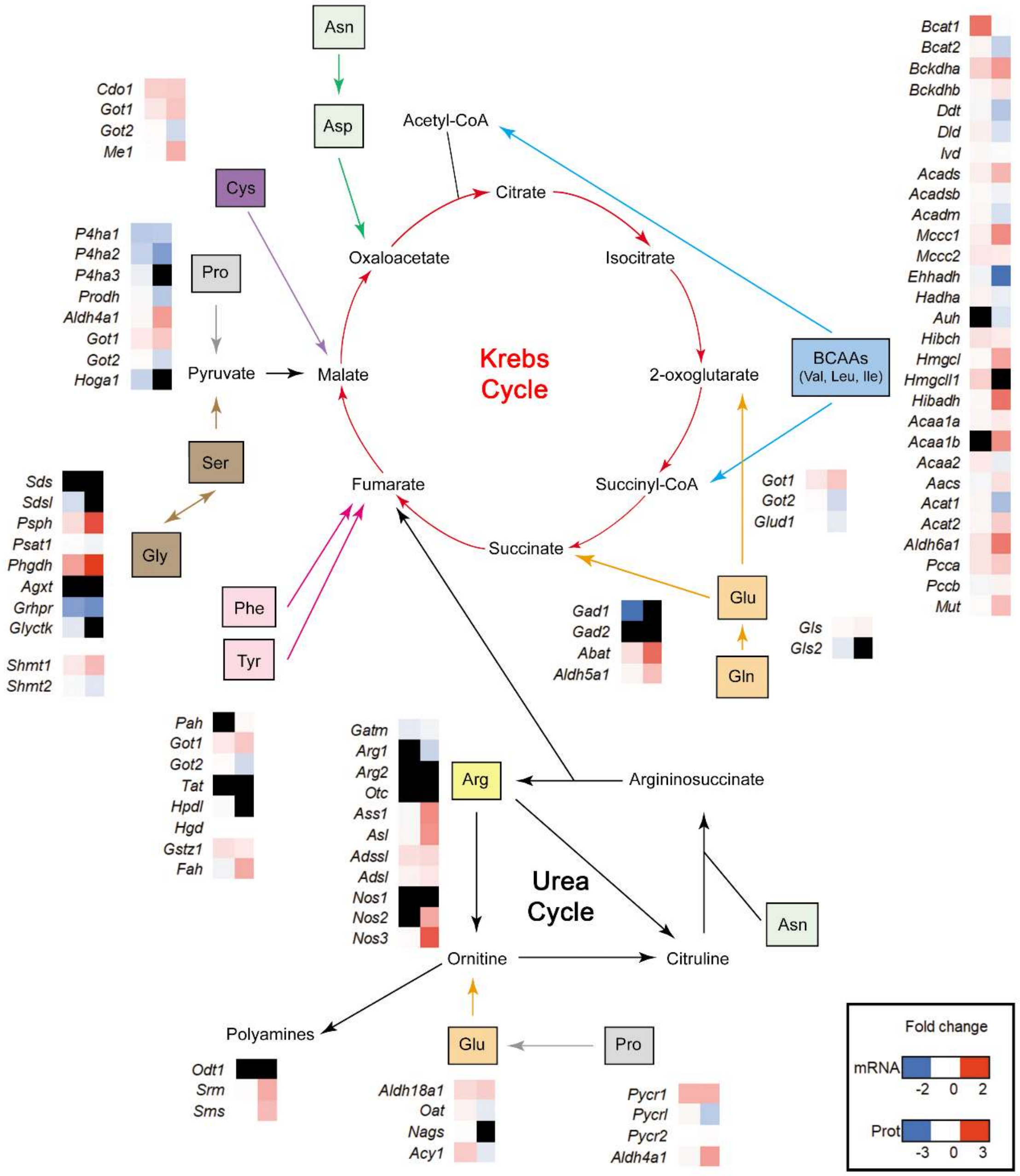


**Figure S5. Schematic representation of amino acid contributions to Krebs and Urea Cycles.** Schematic overview of the transcriptomics (mRNA expression, left column) and proteomics data (standardized protein quantifications, right column) of the re-wired metabolic pathways in the heart of *Hif1a/Nkx2.5* mutants (*Hif1a*$^{f/f}$*/Nkx2.5*$^{Cre/+}$) over control embryos (*Hif1a*$^{f/f}$*/Nkx2*$^{r+/+}$) at E12.5. Data are represented as individual heat maps for the transcript/protein of each pathway calculated as logarithmic Fold Change (logFC) and coded by color intensity following the scale at the bottom. ND indicates no detection. All 14406 expressed genes and 4276 quantified proteins were considered.

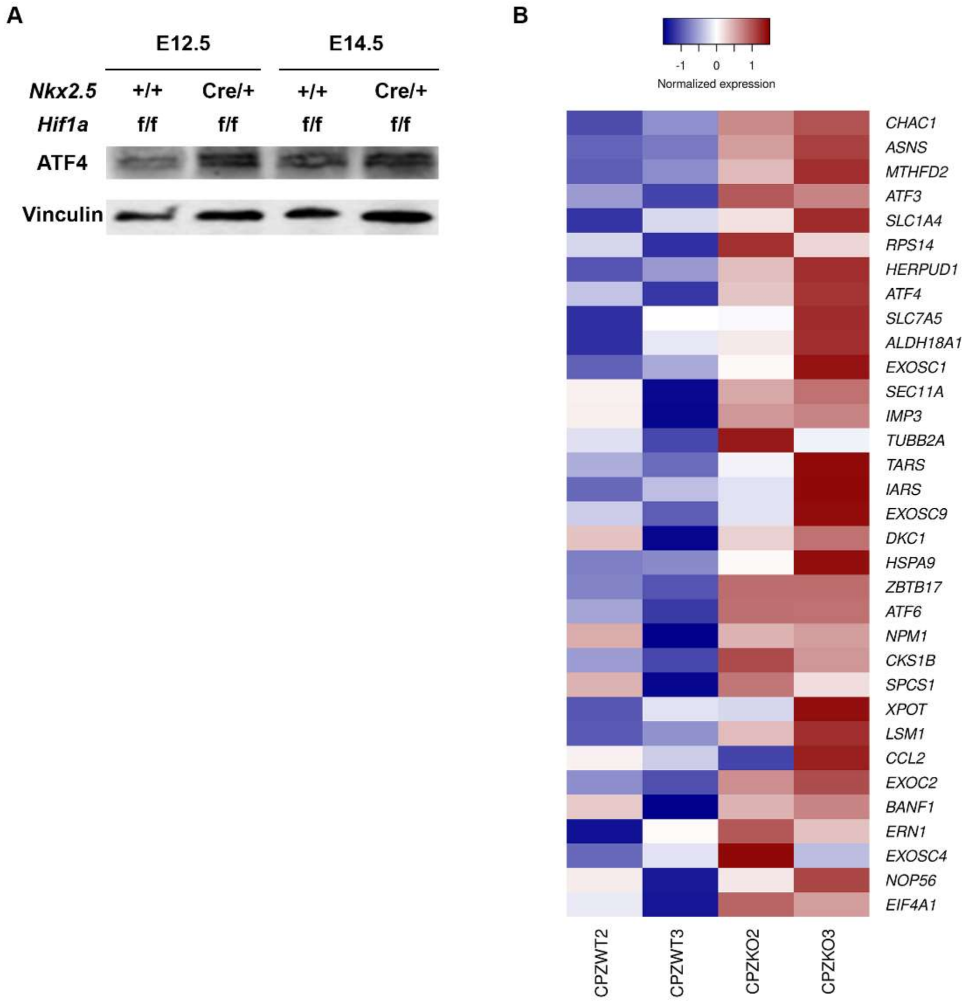


**Figure S6. Induction of ATF4 and other genes involved in UPR upon *Hif1a* deletion. A)** Representative immunoblot against ATF4 (up) and Vinculin (down) in heart lysates of control (*Hif1a*$^{f/f}$/*Nkx2.5*$^{+/+}$) and *Hif1a*/*Nkx2.5* (*Hif1a*$^{f/f}$/*Nkx2.5*$^{Cre/+}$) embryos at E12.5 and E14.5. **B)** Heatmap representing RNA-Seq based, normalized expression levels for genes involved in the Unfolded Protein Response (UPR). The UPR gene set, as defined in the Hallmark database, was detected as enriched in mutant embryos after GSEA, although enrichment was not statistically significant (nominal P value = 0.31). Genes presented in the heatmap correspond to the leading edge subset, this is, those mostly contributing to the calculated enrichment score.

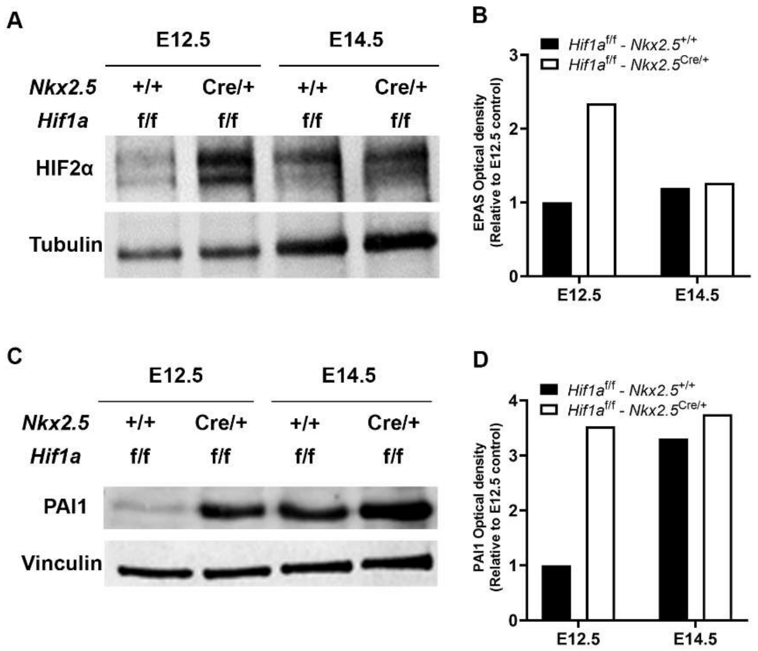


**Figure S7. HIF2α signaling is induced at E12.5 upon *Hif1a* deletion. A)** Representative immunoblot against HIF2α (up) and Tubulin (down) in heart lysates of control (*Hif1a*$^{f/f}$/*Nkx2.5*$^{+/+}$) and *Hif1a*/*Nkx2.5* mutant (*Hif1a*$^{f/f}$/*Nkx2.5*$^{Cre/+}$) embryos at E12.5 and E14.5. **B)** Quantification of band intensity normalized by loading control from immunoblot in A. **C)** Representative immunoblot against PAI1 (up) and Vinculin (down) in heart lysates of control (*Hif1a*$^{f/f}$/*Nkx2.5*$^{+/+}$) and *Hif1a*/*Nkx2.5* mutant (*Hif1a*$^{f/f}$/*Nkx2.5*$^{Cre/+}$) embryos at E12.5 and E14.5. **D)** Quantification of band intensity normalized by loading control from immunoblot in D.
